# Growth anisotropy of the extracellular matrix drives mechanics in a developing organ

**DOI:** 10.1101/2022.07.19.500615

**Authors:** Stefan Harmansa, Alexander Erlich, Christophe Eloy, Giuseppe Zurlo, Thomas Lecuit

## Abstract

The final size and shape of organs results from volume expansion by growth and shape changes by contractility. Complex morphologies arise from differences in growth rate between tissues. We address here how differential growth drives epithelial thickening and doming during the morphogenesis of the growing *Drosophila* wing imaginal disc. We report that 3D morphology results from elastic deformation due to differential growth between the epithelial cell layer and its enveloping extracellular matrix (ECM). Furthermore, the ECM envelope exhibits differential growth anisotropy (i.e. anisotropic expansion in 3D), growing in-plane on one side, but out of plane on the other side. The elasticity, anisotropy and morphogenesis is fully captured by a mechanical bilayer model. Moreover, differential expression of the Matrix metalloproteinase MMP2 controls growth anisotropy of the two ECM layers. This study shows that the ECM is a controllable mechanical constraint whose intrinsic growth anisotropy directs tissue morphogenesis in a developing organ.

## Introduction

During animal development, tissues are shaped by mechanical forces (Gillard and Röper, 2020; Hannezo and Heisenberg, 2019; Vignes et al., 2022). While active contractile forces generated by the cellular actomyosin network have gained substantial attention, the role of cellular volume growth in shaping epithelial tissues as an active process remains less explored (LeGoff and Lecuit, 2016; Stooke-Vaughan and Campàs, 2018). During development, epithelial tissues drastically grow in size due to cell growth (cellular increase in mass and volume) and proliferation (increase in cell number) (Ginzberg et al., 2015). Importantly, both processes contribute to organ growth. The growth rate measures an increase in tissue mass and volume over a time interval. Tissue growth rate may be different within different regions of a tissue or between tissues, a situation which we refer to as differential growth. Differential growth leads to a geometric incompatibility (Aharoni et al., 2016; Eckart, 1948; Lee et al., 2021; Skalak et al., 1997; Truskinovsky and Zurlo, 2019) which is the source of residual stress in the tissue, i.e. the stress that remains in the absence of external forces.

In multilayered tissues such as in organs, differential growth between adjacent layers can lead to mechanical stress that shapes tissues in 3D. For example, differential growth between connected cell layers drives the folding of the human cortex (Garcia et al., 2018; Tallinen et al., 2016), the looping of the gut and formation of villi in the chick (Ben Amar and Jia, 2013; Savin et al., 2011; Shyer et al., 2013), morphogenesis of the airways (Varner et al., 2015) and the heart (Shi et al., 2014).

Here, we use the *Drosophila* wing imaginal disc, a multilayered epithelial structure to study how growth affects morphogenesis of the wing primordium. During larval development, the wing disc grows exponentially. In the presumptive wing territory, called the pouch region, the apical cell surface is spatially organized with larger cell areas in the periphery and smaller in the center (LeGoff et al., 2013; Mao et al., 2013). This gradient of apical cell area was interpreted as a buildup of compressive stress in the center of the disc. Previous studies simplified the disc as a 2D sheet and proposed that differential growth in the plane of the disc epithelium (Mao et al., 2013) as a possible mechanism to explain the observed cell shape gradients.

However, the wing disc is a 3D organ that consists of different layers of material stacked on top of each other, forming a multi-layered sandwich structure. The disc is composed of two stacked epithelial mono-layers: the bottom pseudostratified disc proper epithelium (DP) and the overlying squamous peripodial epithelium (PPE, see Fig.1A). The basal side of each monolayer of this epithelial ‘sandwich’ is surrounded by an extracellular matrix (ECM), effectively making the wing disc a four-layer structure. In particular the ECM has recently gained more attention and was shown to be required for controlling growth (Ma et al., 2017) and morphology of the disc (Atzeni et al., 2019; Nematbakhsh et al., 2020; Sui et al., 2018). Several recent studies investigated the formation of folds surrounding the central portion of the DP, the wing pouch (Sui et al., 2018; Sui and Dahmann, 2020; Tozluoğlu et al., 2019).

**Figure 1.**
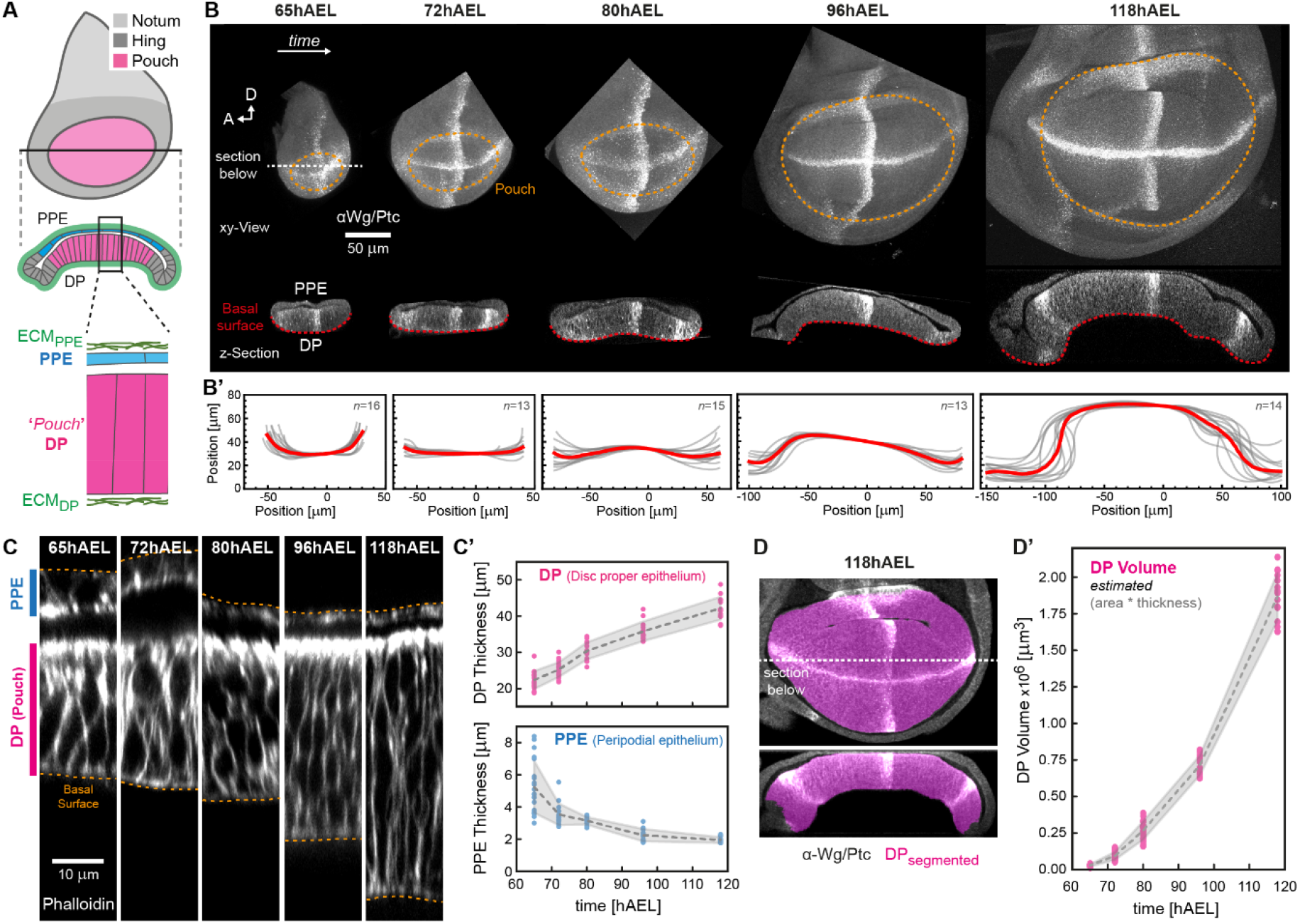
Growth associated epithelial doming and thickening. (**A**) Scheme of the *Drosophila* wing disc: The disc proper epithelium (DP) is overlaid by the peripodial epithelium (PPE, blue). The DP contains the wing pouch (magenta), surrounded by the hinge (dark grey). The disc is covered by an ECM shell (green). (**B**) Wing discs of indicated age in plane (*top*) and section view (*bottom*) stained for Wingless and Patched (Wg/Ptc) to provide landmarks and the tissue outline. Discs in-plane view are oriented with anterior (A) facing right and dorsal (D) facing up, the pouch is outlined by a dashed line. In section view the PPE is facing upwards. The basal surface of the DP is marked (red dotted line). (**B’**) Quantification of average basal surface shape (as indicated by the dotted red line in (B)). Individual outlines are shown in grey, the average outline in red. (**C**) Section views of wing discs (at the A/P-D/V boundary intersection) at indicated age classes. Cell outlines are marked by Phalloidin (F-Actin). (**C’**) Quantification of PPE and DP thickness (measured in sections as shown in C). (**D**) A 118hAEL wing disc marked for Wg/Ptc (grey) after segmentation of the DP pouch volume (magenta) in-plane (*top*) and cross-section view (*bottom*). (**D’**) Quantification of pouch volume (see Fig.S1 and methods for details). Error bands indicate standard deviation.

The ECM is required for cell polarization during early stages of organ formation, especially when lumens are formed (Walma and Yamada, 2020; Zhang et al., 2020). It plays an essential role in organ development, such as during branching morphogenesis (Nerger et al., 2021; Wang et al., 2017). The ECM provides a mechanical and geometric scaffold for the formation of organoids (Nikolaev et al., 2020, reviewed in Hofer and Lutolf, 2021) and embryonic structures in stem cell derived systems (Veenvliet et al., 2020).

How the wing disc acquires its dome-like shape at the end of larval development and how growth and mechanics impact this process remain unclear. Here, we directly assess how the 3D growth properties of the tissue and ECM layers lead to elastic deformation of the wing primordium.

## Results

### Disc curvature increases during development

In order to study the 3D morphology of the growing wing disc we focused on the major growth phase from 65 hours after egg laying (hAEL, mid second instar) to the end of larval development (corresponding to 120hAEL at 25°C). We assessed 3D morphology in crosssections parallel to the anterior/posterior (A/P) axis. While at 65hAEL the basal surface of the DP curves upwards, it becomes flat around 72hAEL and then starts to form an inverse, dome-like structure after 80hAEL (Fig.1B bottom and B’). Coinciding with tissue doming, the thickness of the DP epithelium doubles between 65 and 118hAEL. In contrast, the thickness of the PPE decreases from ~5μm to ~2μm during the same period of development (Fig.1C). Associated with these morphological changes we observe a striking increase in tissue volume. We measured the temporal changes in volume of the central portion of the DP, the wing pouch, and found a ~66-fold volume increase between 65-118hAEL (see Fig.1D, Fig.S1 and methods for details).

Consistent with the tissue scale thickening of the DP epithelium we observed that DP cells become thinner and elongate along their apical-basal axis. The inverse is observed for peripodial cells, which undergo a flattening towards the end of larval development (Fig.S2A-C).

Together, our data show a concomitant increase in tissue size with thickening and bending of the disc epithelium. This raises the question of how growth and tissue morphology are linked during wing disc morphogenesis.

### Tissue thickening is not due to differential cell growth in the tissue plane

We first considered the possibility that the doubling in tissue thickness stems from spatially non-uniform growth in the plane of the DP tissue. Previous work, using indirect measures of growth such as cell counts in clones, proposed that cell proliferation is increased in the center versus the periphery of the disc, leading to stress accumulation in the disc center (Mao et al., 2013). Consistently, artificially increasing growth in clones of cells leads to the accumulation of stress in neighboring cells but not to an increase in thickness (LeGoff et al., 2013; Pan et al., 2016).

To test theoretically if increased central growth can drive tissue thickening, we constructed a model based on the conceptual framework of morphoelasticity (Rodriguez et al., 1994; Ambrosi and Guana, 2005; Goriely, 2017) using finite element simulations (Hosseini et al., 2014; Taber, 2008). We model the DP tissue layer as an elastic continuum object, which can be pre-stressed through a growth tensor field 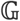. 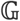 determines the local addition of mass and how the geometry is modified due to growth. Thus, by controlling 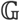 we can model that more volume is added towards the center of the disc than in the periphery, and computationally explore the effects on tissue shape and stress (see SI for details). The simulations indicate that increased growth in the DP center versus the periphery can induce tissue thickening, although a 2-fold growth mismatch generates at most a 16% increase in tissue thickness (Fig.2A-B), while experimental data indicate that the DP tissue thickness doubles.

**Figure 2.**
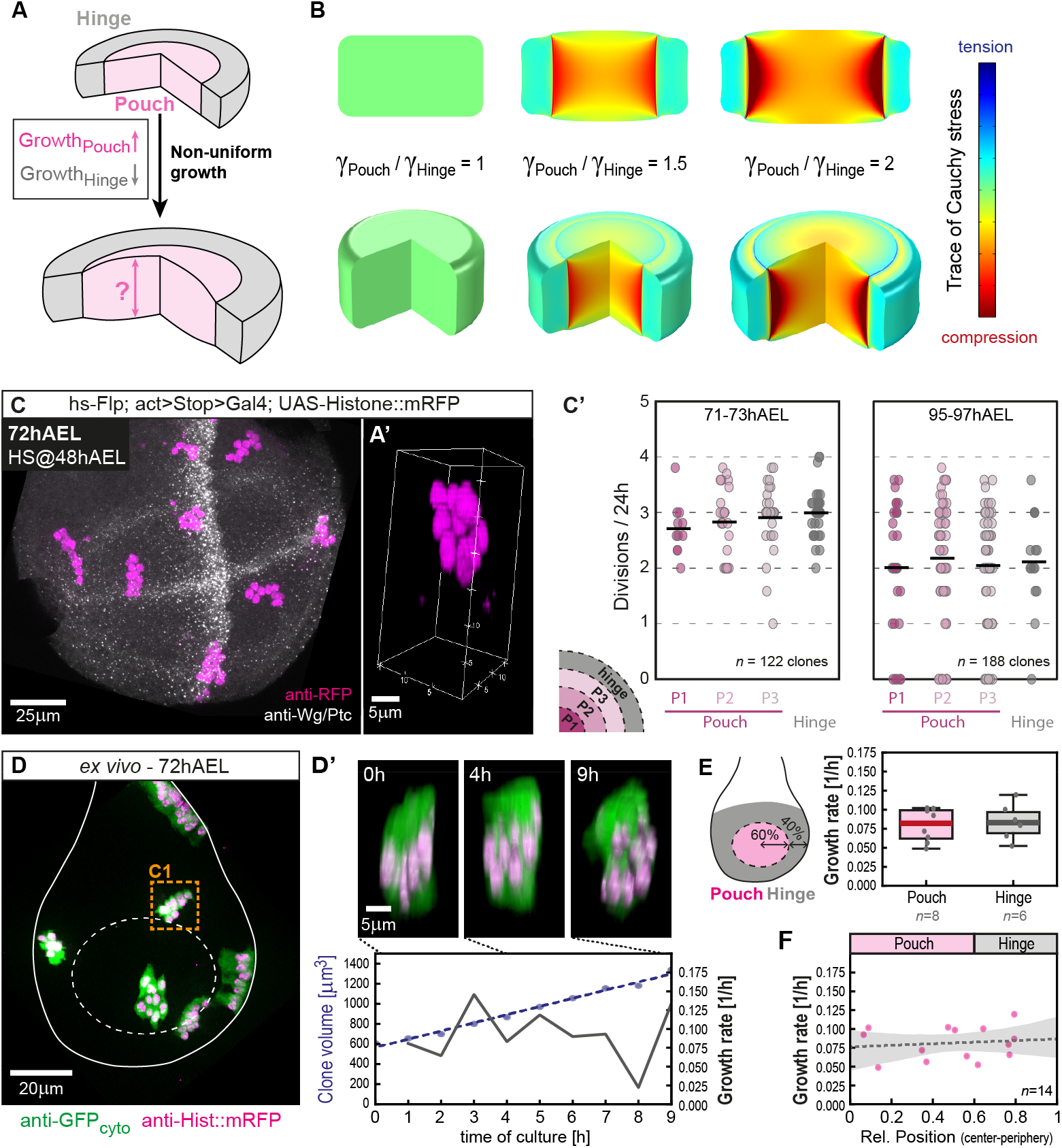
Growth is homogeneous in the plane of the DP epithelium. (**A**) Growth non-uniformities with higher growth in the DP (magenta) versus the surrounding hinge (gray) tissue can result in epithelial thickening. (**B**) Results of simulations for different ratios of the growth factor γ between central Pouch (γ_Pouch_) and peripheral Hinge (γ_Hinge_). (**C**) Representative 72hAEL wing disc expressing Histone::RFP in a clonal manner (24h after clone induction). 3D-volume view of an example clone is shown on the right. (**C’**). Clonal division rates at 72hAEL (*left*) and at 96hAEL (*right*) are assessed in 4 elliptic domains covering the pouch (P1-P3, central to peripheral) and the hinge. (**D**) *ex vivo* cultured 72hAEL wing disc expressing cytosolic GFP (clone volume) and Histone::RFP (nuclei) in a clonal manner at the beginning of culture. The clone marked by the orange rectangle is shown in 3D-view in (D’) at indicated times of culture. *bottom*: Clonal volume and growth rate over 9h of culture for the shown clone. (**E**) *left*: Clones were classified as ‘pouch’ or ‘hinge’, depending on their position. *right*: Average growth rate in the pouch (magenta) and in the hinge region (gray) at 72hAEL. (**F**) Data shown in (E) plotted in respect to relative clonal position (center to periphery). A linear regression is shown by a gray dashed line (error band indicates standard deviation).

We assessed cell growth extensively using 3 independent methods. We first investigated the spatial pattern of cell divisions by clonal assays (as done in (Mao et al., 2013)), making sure that clonal density is low (on average 9 clones/disc at 72hAEL, see methods) and hence fusion unlikely to occur. We found no evidence of differential growth between the central pouch and peripheral hinge region from 48-96hAEL (Fig.2C). We next stained staged and fixed samples for Phospho-Histone H3, a marker for mitosis (Fig.S3A), and observed a uniform pattern of proliferation density from 65hAEL to the end of larval growth.

Importantly, growth is a dynamic and 3-dimensional process. We therefore established volumetric 3D live-imaging in *ex vivo* culture of wing discs to directly measure clonal growth rates (see methods). We have followed volumetric changes in small clones containing few cells for up to 10h in disc explants at 72hAEL, the time point when DP thickness starts to increase (Fig.2D). We also found that the cell growth rate was uniform (0.085 h^−1^) within the plane of the DP epithelium (Fig.2E-F).

In conclusion, our results indicate that growth is uniform in the plane of the DP epithelium at different stages of larval development while simulations suggest that a large difference in growth rates are required to induce significant thickening. Therefore, tissue thickening has a different origin than differential cell growth in the plane of the DP epithelium.

### Bending is not due to differential growth of the tissue layers

Next, we considered the 3D multilayered structure of the disc consisting of two growing tissue layers: the DP and the PPE. We used the concept of morphoelasticity to gain insight into this growing bilayer structure. Using a non-linear elastic continuum model we simulated two mechanically coupled and growing elastic layers (DP and PPE, see Fig. 3A). Morphoelasticity considers that during growth, changes in tissue shape result from both an increase in volume and elastic deformations. The change in tissue shape (the total deformation) is described by the deformation gradient tensor 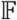. Indeed, 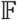 captures the transformation of a growing structure from its original stress-free state (early in development), called the initial state, to its grown, final shape and size (observed state, Fig.3A). Importantly, this transformation is due to a growth component, described by a growth deformation tensor 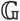, and an elastic component, described by the elastic deformation gradient tensor 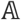 (such that 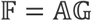).

**Figure 3.**
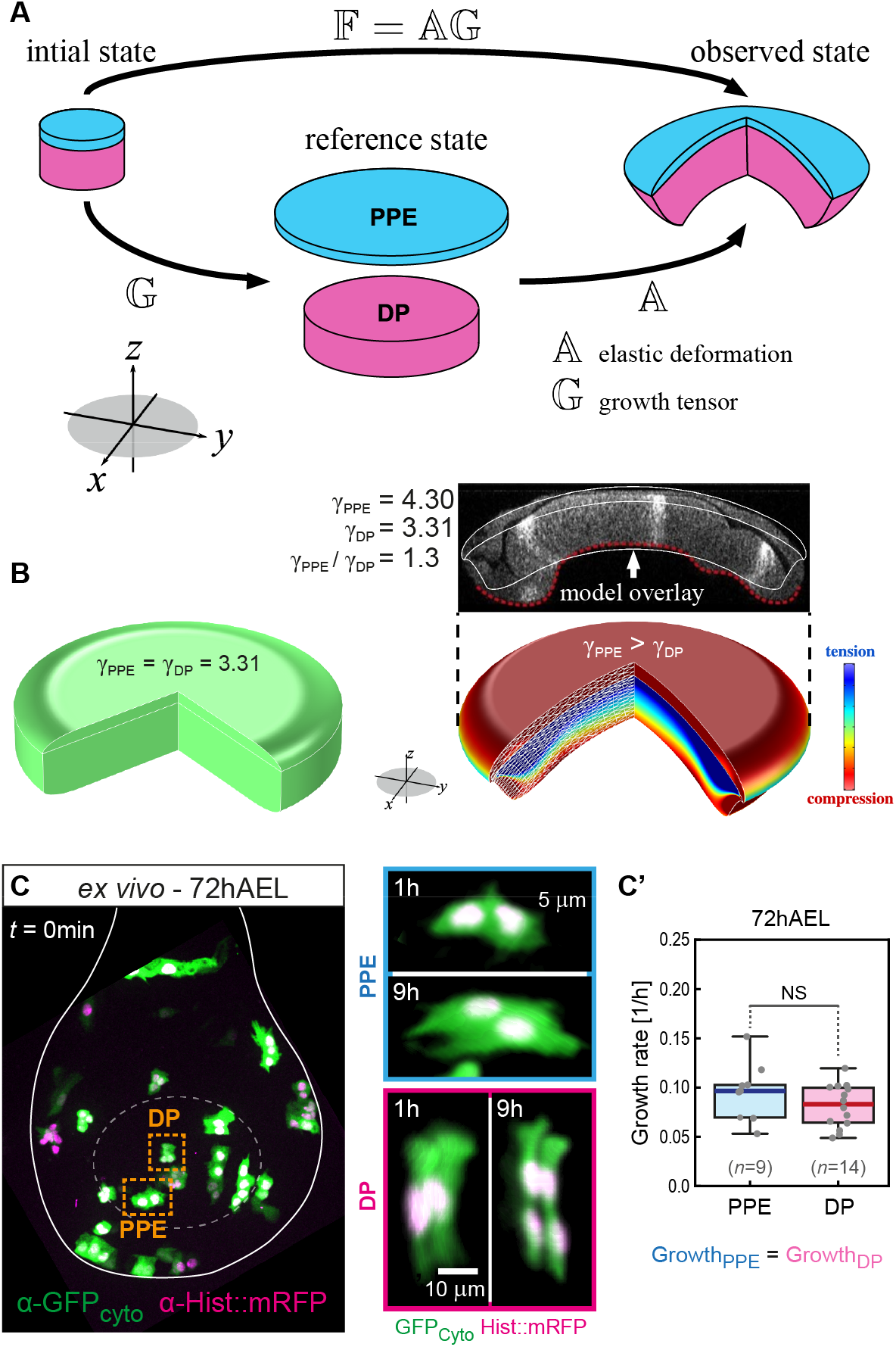
Non-uniform growth between epithelial layers does not drive tissue bending. (**A**) Geometric decomposition of the wing disc as a sandwich of growing elastic layers, the PPE (blue) and the DP (magenta), each growing uniformly in plane. The growth tensor 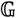 describes growth without stress; different growth rates in the respective layers (described by 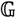) lead to different sizes of the individual discs. Elastic deformation (described by 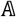) connects the discs into one coherent object, leading to the observed morphology and accumulation of residual stress. (**B**) *left*: If the growth rates in all layers are equal, growth is compatible, leading to no residual stress (green color). *right*: If the PPE volume grows ~70% times faster than the DP, simulations approximate well the curvature and morphology of 118hAEL wing discs (see inset with model overlay), with the PPE being in compression (red). (**C**) *left*: Maximum projection of an *ex vivo* cultured 72hAEL wing disc at the start of imaging (0min) containing clones of cytosolic GFP (volume) and Histone::RFP. *right*: 3D projection of two example clones at the beginning and end of culture. (C’) Average clonal growth rates at 72hAEL.

We assume that in the initial state, the two tissue layers have the same disc size and can be connected into a stress-free structure (Fig.3A). The growth tensors 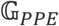 and 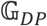 describe the growth of the PPE and the DP layers, respectively. When growth between the PPE and DP layer differs, namely 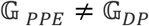, the two grown tissue layers DP and PPE have different sizes that no longer form a coherent body, which is called a geometric incompatibility (see reference state, Fig.3A). However, the two different sized discs can be connected into one coherent object by elastic deformation (by 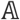) which builds residual stress. Therefore, the observed state is residually stressed and elastically deformed (Fig.3A). The assumption that the wing disc is an elastic material with extensive residual stress is supported by the fact that cutting the domed wing disc in 3^rd^ larval stage, leads to nearly instantaneous relaxation of the shape (Fig. S3B). This is also further confirmed later. We describe the material of the layers by a nearly incompressible neo-Hookean material, a commonly used material model for morphogenetic tissue due to its relative simplicity and ability to describe large deformations (Hosseini et al., 2014; Pence and Gou, 2015; Taber, 2008). In a polar cylindrical basis 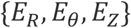, we denote components of the growth tensor 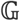 by 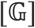 in the DP and PPE as

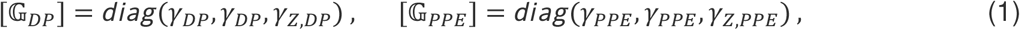

respectively. Here, *γ_DP_, γ_PPE_* are the in-plane growth parameters in the respective layers (the plane direction is marked as a gray circle in the coordinate system in Fig. 2A). The growth parameters in the axial, or *Z*-direction which is orthogonal to the plane are denoted *γ_Z,DP_, γ_Z,PPE_*. When *γ_DP_* = *γ_Z,DP_*, then growth is isotropic, i.e. the same in all directions. In the case *γ_DP_* ≠ *γ_Z,DP_*, growth is anisotropic. A subcase of anisotropic growth is planar growth, where *γ_Z,DP_* = 1.

We assume planar growth for the two tissue layers, since in epithelia the cell division plane is parallel to the epithelial plane (van Leen et al., 2020) (*γ_Z,PPE_* = *γ_Z,DP_* = 1). First, we considered a scenario where the two layers grow equally (*γ_PPE_* = *γ_DP_*, compatible growth) and no elastic stress and no bending is produced through growth (Fig.2B, *left*). In a second scenario, when the PPE grows faster than the DP layer (*γ_PPE_* > *γ_DP_*), we find that qualitatively correct bending is predicted (Fig.2B *right*). In order to obtain bending that is comparable to wing discs at 118hAEL the model predicts that the PPE layer grows ~70% more in volume than the DP layer (see overlay, Fig.2B *top*). Moreover, the PPE layer is compressed (red color) in this scenario. In order to experimentally test this prediction we measured growth rates in DP and PPE clones in *ex vivo* cultured discs (Fig.2C). However, at the time point of bending onset (72hAEL) clonal growth rates over a period of 9 hours are comparable in both epithelial layers. To further test the mechanical role of the PPE layer, we reduced growth in peripodial cells by overexpression of a dominant-negative form of Phosphatidylinositol 3-kinase (PI3K_DN_), a major growth regulator. In this situation, PPE growth is significantly reduced, but bending of the DP not altered (Fig.S3D-G).

In summary, these results demonstrate that epithelial doming and thickening are not due to a non-uniformity of growth within or between epithelial layers. Furthermore, given comparable growth rates in the DP and PPE layer, the two layers can be mechanically seen as a single layer due to their compatible growth.

### The ECM is essential for tissue bending

Given that growth in the tissue layers is homogeneous, we next considered the ECM layers. The basal surface of the wing disc is covered by a sheet-like ECM shell that is rich in Collagen IV, called Viking (Vkg) in *Drosophila* (Lunstrum et al., 1988) Given that a loss of Integrin adhesion (Domínguez-Giménez et al., 2007), *vkg* mutations (Pastor-Pareja and Xu, 2011) or the acute loss of the ECM layer (Ma et al., 2017; Nematbakhsh et al., 2020) were shown to impair tissue thickening and doming, we next tested if the ECM acts as a geometric and mechanical constraint during tissue growth.

A continuous layer of ECM, visualized by a GFP-tagged version of Vkg (Vkg::GFP) (Morin et al., 2001), is covering the basal surface of the DP and PPE (Fig.4B, top). Thus the ECM forms an elastic boundary around the wing disc throughout its growth and morphogenesis. In order to investigate the role of the ECM shell, we acutely removed the ECM by Collagenase digestion at different stages of development. At 96hAEL, when the disc is fully domed, this led to an inversion of tissue curvature, consistent with earlier report (Nematbakhsh et al., 2020) (Fig.S4A+B). We reasoned that residual stress due to apical Myosin II (MyoII) contractility would underlie this inverted curvature. Indeed, acute ECM digestion and concomitant inhibition of Myosin II (see methods for details) led to a complete flattening of the epithelial layer from 72hAEL onwards, when disc curvature arises (Fig.4B bottom and B’). At 65hAEL, collagenase treatment left the disc in its flat configuration. Importantly, the inhibition of MyoII alone does not affect overall disc morphology so the ECM plays a primary role. These findings show that MyoII is negligible, but the ECM layer is essential for wing disc bending from late 2^nd^ instar onward.

**Figure 4.**
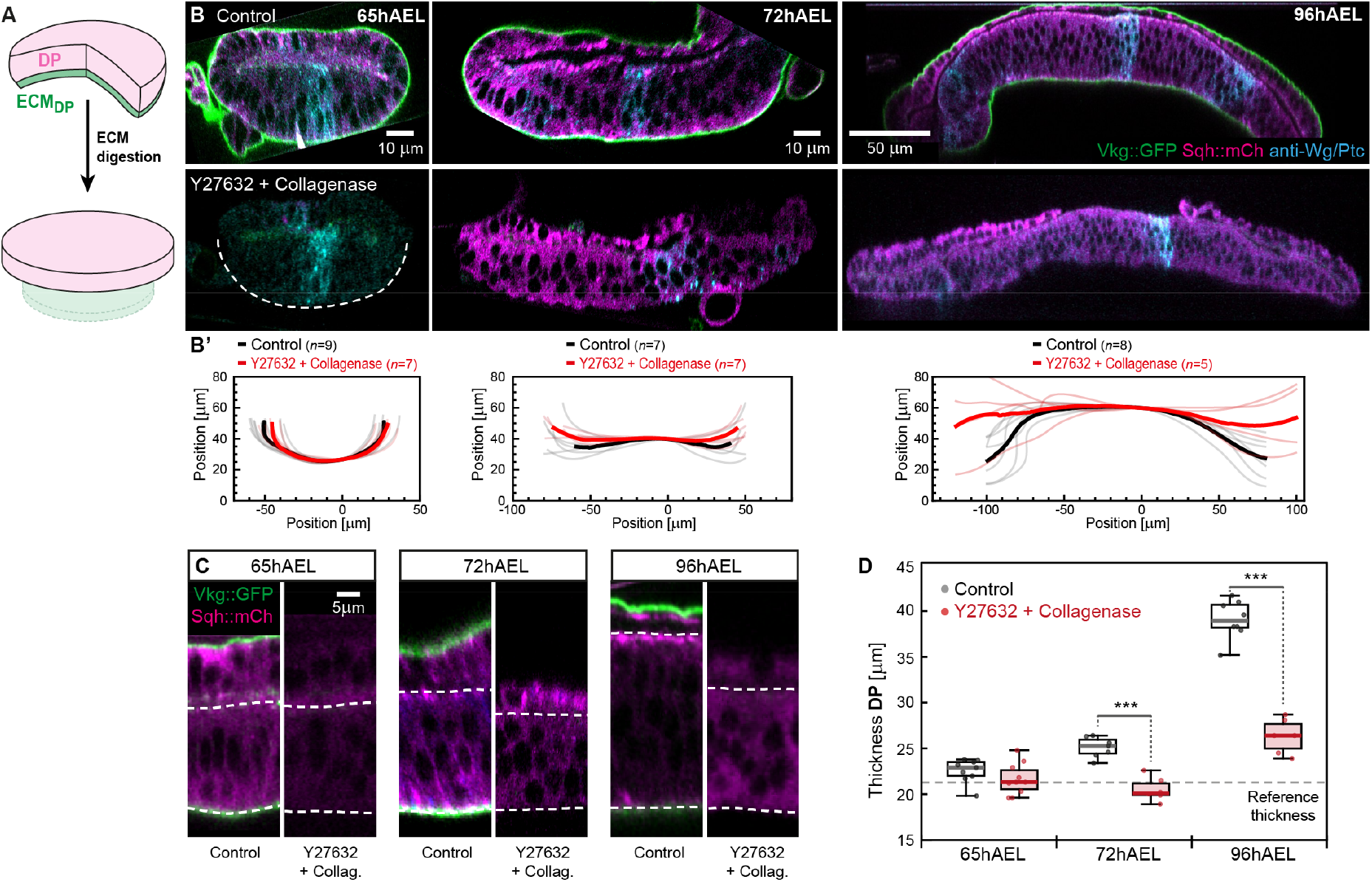
The ECM constraints *planar* epithelial growth and is required for disc doming. (**A**) Acute digestion of the ECM shell by Collagenase leads to relaxation of the epithelial layers. (**B**) *top*: Representative sections of control wing discs at indicated stages of development marked for the ECM (Vkg::GFP, green) and MyoII (Sqh::mCh, magenta). *bottom*: Representative wing discs after acute digestion of the ECM by Collagenase and inhibition of MyoII (by Y27632, see methods). Dashed line marks DP basal outline at 65hAEL. (**B’**) Average basal outlines before (black line) and after ECM digestion + Myosin II inhibition (red line). (**C**) Cross sections of control and ECM digested + MyoII inhibited wing discs. Apical and basal surface of the DP are marked by dashed lines. (**D**) Quantification of epithelial thickness (as seen in (C)) upon digestion of the ECM and inhibition of MyoII (red dots). The reference thickness observed at 65hAEL is indicated by a dashed line.

### Tissue thickening results from in-plane tissue growth and ECM elastic constraints

In addition to the loss of epithelial bending, we also observed a significant decrease of epithelial thickness upon loss of the ECM and MyoII contractility after 72hAEL (Fig.4C+D). At all observed time points the digestion of the ECM returns the epithelial thickness close to the values observed in control discs at 65hAEL. In contrast, at 65hAEL no relaxation upon ECM digestion is observed, suggesting that it can be considered as stress-free reference configuration.

The fact that the relaxed tissue thickness remains nearly constant over time indicates that the DP layer predominantly grows in-plane and that the observed thickness increase is mainly due to elastic compression mediated by the ECM around the whole disc.

### Differential volume growth between DP and associated ECM

Although tissue bending requires an intact ECM layer, the ECM is present on both sides of the disc, facing the DP and PPE tissue layers. It suggests that the ECM may have different properties on both sides of the disc. We considered explicitly the growth of the ECM layers on both sides of the disc and hypothesized in particular that a volumetric growth mismatch between the epithelial and the ECM layer is responsible for the build-up of stress and the observed morphology.

We first investigated the thickness of the ECM, visualized by Vkg::GFP, at defined stages of development (see methods). We find that the thickness of the bottom ECM_DP_ increases by ~36% from 72 to 118hAEL (Fig.S5A+D). In contrast, the thickness of the top ECM_PPE_ layer does not significantly change during this time window (Fig.S5B).

We quantified the growth of the ECM_DP_ covering the basal surface of the central portion of the DP, the wing pouch, following two approaches. First, we estimated ECM_DP_ volume by multiplying ECM thickness with the respective area values of the pouch (see methods for details). Secondly, we measured the increase in integrated fluorescence intensity of the Vkg::GFP signal underlying the pouch (marked by the Wg ring) between 65 and 118hAEL (Fig.S5F). When we compared the relative increase in pouch volume with the relative increase in ECM estimated volume or integrated intensity we found that the pouch outgrows the ECM_DP_ by ~15-20% (Fig.S5E+G). Notably, we found a similar volumetric increase for the PPE layer and its associated ECM_PPE_ layer (~7% difference, see Fig.S5C).

In summary, these findings show that the DP tissue layer outgrows its ECM_DP_ layer, leading to an effective volumetric growth mismatch between the two layers. Modeling this 20% growth mismatch was however not sufficient to predict correct tissue geometry (See Fig. S5H). In contrast, the PPE and its ECM_PPE_ layer show similar volumetric growth suggesting compatible growth of these two layers. This supports our hypothesis that the bottom ECM_DP_ acts as a geometric constraint for epithelial growth.

### Spatial differences in ECM growth anisotropy

So far we considered the difference in volume increase of the ECM versus the overlying DP tissue layers. However, growth is described by a tensor so we next considered growth anisotropy. Specifically we addressed whether there is a difference in growth anisotropy between the ECM layers (*γ_Z,ECM_* ≠ 1) and the tissue layers which, as we showed (Fig. 4) grow in a plane (*γ_Z,DP_* = 1). We hypothesized that the top ECM_PPE_ and bottom ECM_DP_ layers have different growth anisotropy.

In order to assess differences in ECM growth anisotropy we aimed to eliminate elastic deformation (due to epithelial growth) and visualize the relaxed ECM layers in their unstressed configuration (‘reference state’). This was achieved by exposing disc explants to the detergent Triton X-100 (referred to as ‘decellularization’ in the following, see methods) which results in the degradation of the lipid bilayer and a loss of cells and thus hydrostatic pressure on the surrounding ECM (Fig.5A+B). Chemical decellularization was used in various regenerative (Chen et al., 2016; Garcia-Puig et al., 2019; Sonpho et al., 2021) and biomedical approaches (Neishabouri et al., 2022) and decellularization by Triton X-100 retains ECM microstructure (Fernández-Pérez and Ahearne, 2019). The decellularization of 118hAEL discs results in a degradation and shrinking of the epithelial layers, well visible in cross-section views (Fig.5B *top* versus *bottom*).

**Figure 5.**
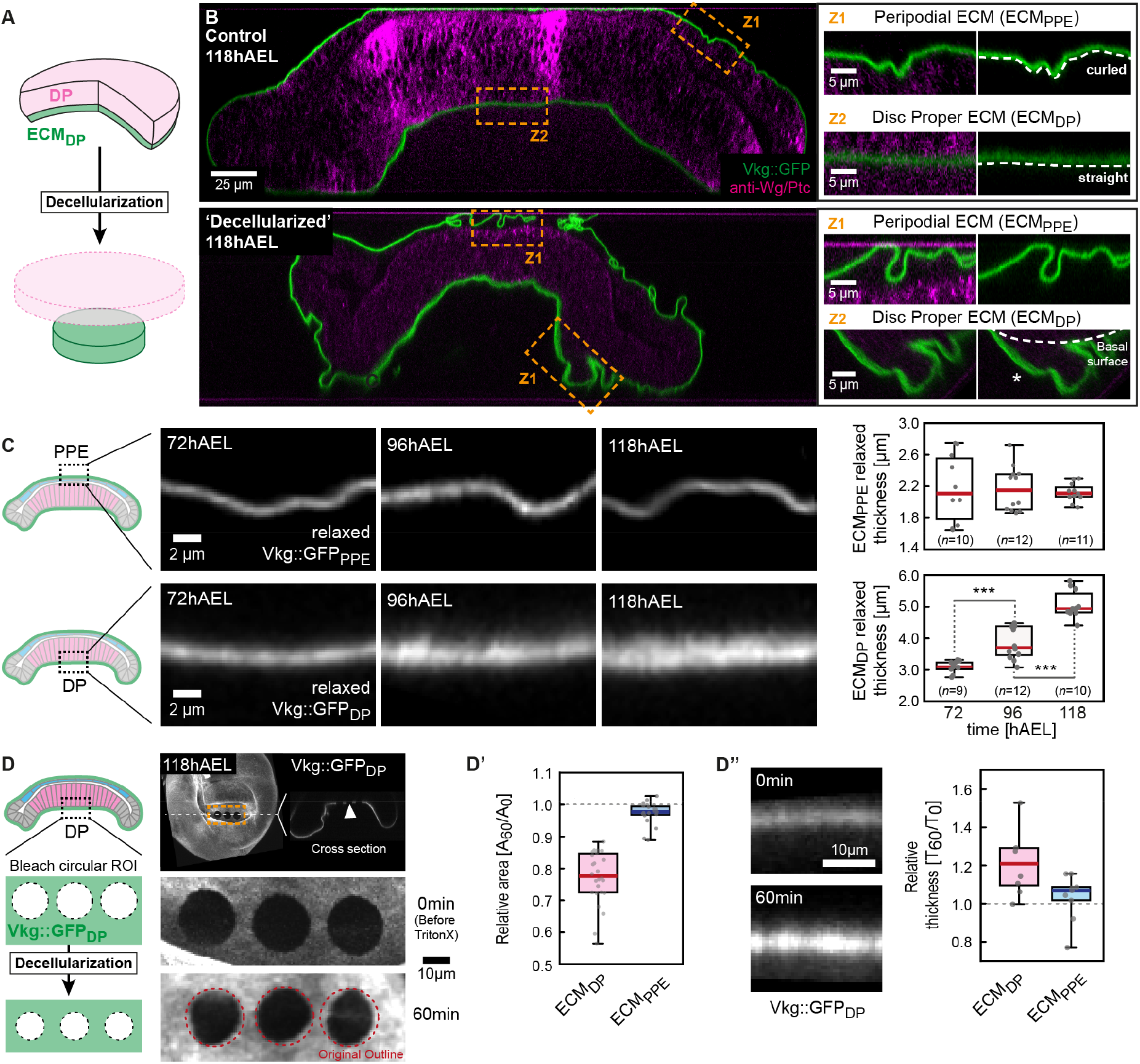
Spatial differences in ECM growth anisotropy amplify stress accumulation and allow symmetry breaking. (**A**) Acute removal of cellular pressure by decellularization reveals the relaxed configuration of the ECM. (**B**) Cross sections of a control wing disc (*top*) and a wing disc after decellularization (*bottom*, see methods). The ECM is marked by *Vkg::GFP* (green) and the epithelial layers by a staining for Wg/Ptc (magenta). *right*: Magnifications of indicated regions. The relaxed geometry of the ECM was assessed in regions where the ECM had separated from the epithelial layer (see asterisk). (**C**) Cross-sections of relaxed sections of the top ECM_PPE_ (marked by Vkg::GFP_PPE_, *top*) and the bottom ECM_DP_ (Vkg::GFP_DP_, *bottom*) at indicated stages. *right*: Quantification of relaxed ECM thickness (see methods for details). (**D**) Circular regions of interest (ROI) were bleached on the ECM Vkg::GFP signal (see scheme on the left). Upon decellularization, changes in circular area (D’) and ECM thickness (D”) were quantified in the top ECM_DP_ and bottom ECM_PPE_ layer.

We first investigated changes in relaxed ECM thickness upon decellularization at defined stages of development (72, 96 and 118hAEL). Strikingly, the relaxed thickness of the top ECM_PPE_ did not change during larval development (Fig.5C top and Fig.S6A), while the relaxed thickness of the bottom ECM_DP_ increases by ~40% between 72 and 118hAEL (Fig.5C bottom and Fig.6B). Therefore, the top ECM_PPE_ layer follows planar growth (*γ_Z_* = 1) that does not result in any thickness change over time. In contrast, the growth of the bottom ECM_DP_ is markedly non-planar (*γ_Z_* ≠ 1), as it grows in thickness as well as in plane.

**Figure 6.**
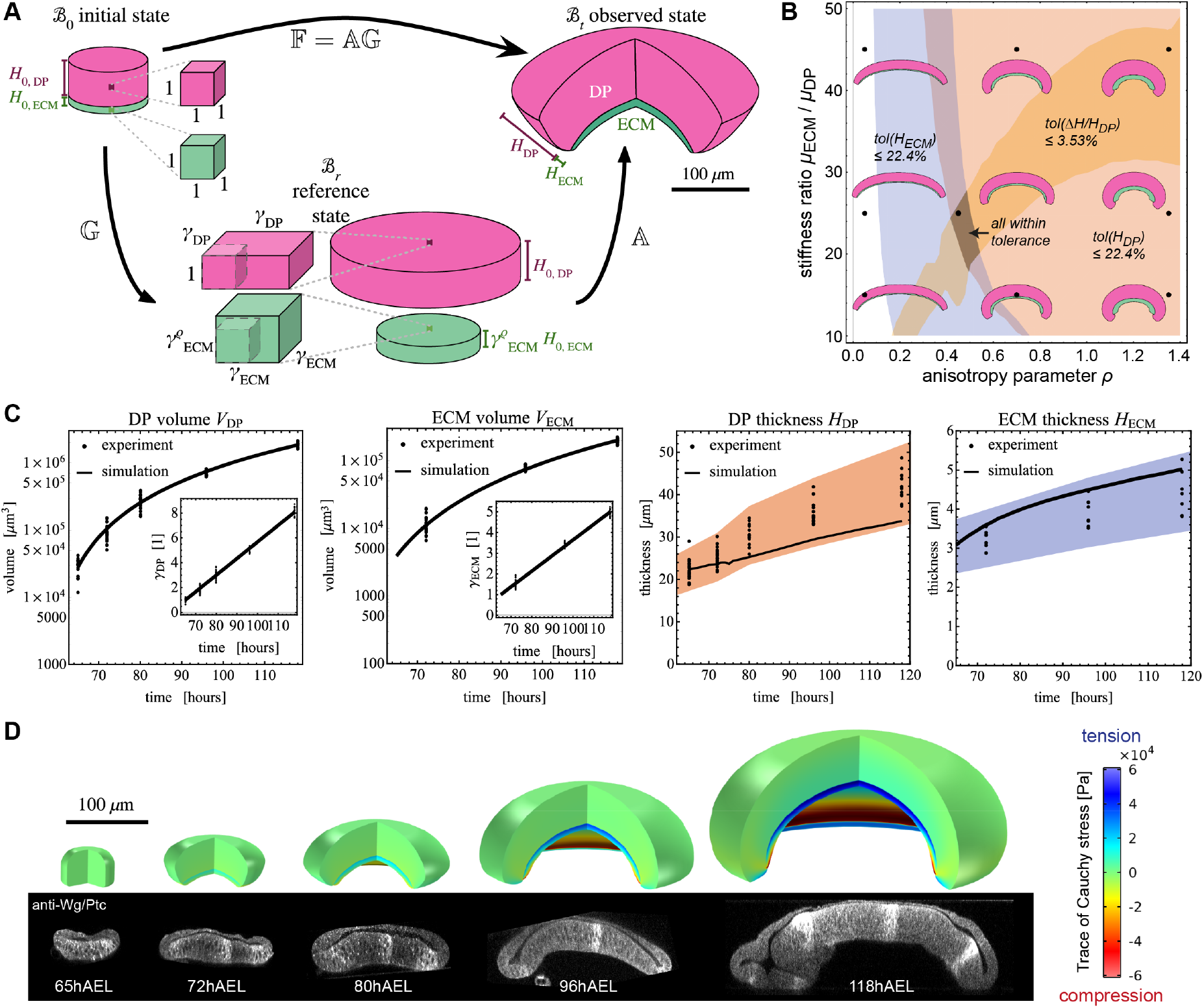
An elastic bilayer model captures growth induced epithelial morphogenesis. (**A**) Geometric decomposition for a bilayer structure, composed of DP (purple) and ECM_DP_ (green). In the initial state 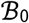, the disc is unstressed and undeformed. Representative volume elements with unit lateral lengths are shown. The DP and ECM thickness is denoted *H*_0, *DP*_ and *H*_0, *ECM*_, respectively. The growth tensor 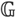 describes the transformation to the reference state 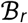 which is grown and relaxed. Since the DP grows in plane, its thickness remains *H*_0, *DP*_. The ECM grows orthogonally to the plane as controlled by the anisotropy parameter *ρ*, and the relaxed thickness is greater than *H*_0, *ECM*_. Through the elastic deformation gradient 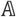, stress is introduced, leading to the domed observed state 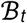. (**B**) Parameter diagram to determine a region in which simulation values are within chosen tolerances of experimental measurements. In the blue, red and orange regions, the simulation results are within tolerance of the measured ECM thickness *H_ECM_*, DP thickness *H_DP_*, and relative increase in thickness upon decellularization (see SI for details). In the dark region, all three conditions are satisfied simultaneously. Parameters *ρ* = 0.45 and *μ* = 25 were chosen as the best fit, of which the morphology is shown in the central out of 9 insets showing cross-sections of the simulated wing disc. The result *ρ* = 0.45 suggests that the bottom ECM_DP_ growth anisotropy lies midway between planar (*ρ* = 0) and isotropic growth (*ρ* = 1). (**C**) Model predictions (*ρ* = 0.45, *μ* = 25) compared to experimental data. *Left two plots*: The insets show linear profiles of *γ_DP_* and *γ_ECM_*. *Right two plots*: Comparison between simulated and experimental values for DP thickness *H_DP_*, and ECM thickness *H_ECM_*. The colored bands show the region within 22.4% of the mean of experimental values, in which the simulation values are contained. (**D**) The shape of the simulated wing disc, using the best fit parameters, is shown compared to cross-sections of representative wing discs. The simulated discs are shown with a quarter of each disc removed for illustration purposes (simulations are axisymmetric).

To monitor more directly the elastic relaxation of the ECM we decellularized disc explants *ex vivo* and used live-imaging to follow changes in ECM area and thickness. Due to a lack of traceable landmarks in the ECM we bleached circular regions of interest (ROI) on the Vkg::GFP labeled ECM (Fig.5D, see methods). Following decellularization, a previously stretched ROI is expected to relax in area and to thicken concomitantly. In the top ECM_PPE_ neither the circular area nor ECM thickness changed after the ECM_PPE_ reached an equilibrium configuration after 60min (Fig.S6D). This confirms that the upper ECM_PPE_ layer is not under mechanical load and grows compatible with the underlying PPE layer, in the plane. Similar analysis in the bottom ECM_DP_ layer showed that the bleached circular area decreased to ~79% of the original area after one hour (Fig.5D’ and Fig.S6C). Concomitantly, the bottom ECM_DP_ thickness increased by ~25% (Fig.5D’’), confirming that the bottom ECM layer is stretched elastically.

In summary these results demonstrate that the three top layers (ECM_PPE_, PPE and DP) all show compatible planar growth. In contrast, growth of the bottom ECM_DP_ is incompatible with the rest of the disc because it also grows in thickness. Therefore, the 4-layered wing disc can be mechanically simplified by merging the three top layers into one layer. The merged top layer and the bottom ECM_DP_ layer, build stress and curvature through incompatible growth.

### Differential growth anisotropy recapitulates tissue shape changes

In order to test whether such differences in growth anisotropy can predict quantitatively the observed morphogenesis of the disc, we next modified the non-linear elastic bilayer model. Since the PPE and ECM_PPE_ grow compatibly with the DP layer their effect on mechanics and shape can be neglected, and therefore were excluded from further modeling. We consider a structure consisting of the DP layer at the top and the ECM_DP_ layer at the bottom (Fig. 6A). Our hypothesis can thus be stated as the DP tissue following planar growth, and the ECM_DP_ following non-planar growth (*γ_z_* > 1).

In the DP tissue layer growth is planar (see Fig. 4), hence

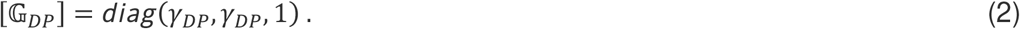

Here, *γ_DP_* was determined through a least-square fit of the experimental volumetric data (see SI).

In the previous section we showed that the thickness of the bottom ECM layer increases with time and hence follows a non-planar mode of growth. Since the extent of growth anisotropy is not known we introduce a positive *growth anisotropy parameter ρ* for the bottom ECM layer:

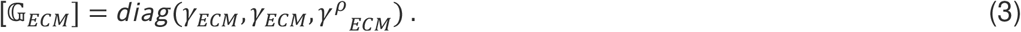

If *ρ* = 0, the ECM would grow in the plane like the DP, see Eq. (2). If *ρ* = 1, the ECM would grow isotropically, like the swelling of a hydrostatic gel (Dolbow et al., 2004; Sultan and Boudaoud, 2008). If *ρ* > 1, growth would be primarily in *Z*-direction, i.e. a form of surface growth by accretion, similarly to the growth of shells and horns (Erlich et al., 2016; Skalak et al., 1997).

Analogously to the DP, we obtained *γ_ECM_* via a linear fit of the experimental volume data, but *γ_ECM_* remains a function of *ρ* (see SI).

Therefore, two dimensionless parameters required to parameterize the model remain to be determined: the growth anisotropy parameter *ρ* and the ratio of elastic moduli, *μ* = *μ_ECM_/μ_DP_*, where *μ_ECM_* and *μ_DP_* are the shear moduli of the respective layers. These parameters *ρ* and *μ* are determined by comparing the model results to three experimentally measured quantities: The disc proper thickness *H_DP_*, the ECM thickness *H_ECM_*, and the relative thickness increase of the ECM upon decellularization. Fig. 6B shows a phase diagram of tissue shape as a function of *ρ* and *μ*. In the gray region, where all colored regions overlap, the experimental and simulated values of all three quantities are within tolerance (see SI). As the best fit from that region, we determined *ρ* = 25 and *μ* = 0.45. The latter result suggests that bottom ECM_DP_ growth anisotropy lies midway between planar (*ρ* = 0) and isotropic growth (*ρ* = 1). Fig. 6C shows the simulated versus experimentally measured values for this best fit (see SI). The thickness values *H_DP_, H_ECM_* are given alongside a tolerance interval that quantifies the relative error between experimental means and simulated values. We can see that the simulations capture well the trends of the increasing thicknesses. The shape of the simulated wing disc (Fig.6D), using the best fit parameters for *ρ* and *μ*, fit well to experimentally obtained morphologies and reveal an increasing buildup of tension in the ECM and compression in the DP.

In summary, a relatively simple mechanical model of growth, incorporating distinct growth anisotropies in the DP and bottom ECM layers, recapitulates wing disc morphogenesis in 3D.

### Mmp2 is required for planar ECM growth

The ECM growth anisotropy is different in the PPE and DP layers. We next addressed the mechanisms controlling planar versus more isotropic ECM growth. We next hypothesized that the difference in growth anisotropy is due to differential expression of ECM modifiers in the PPE versus the DP layer. A potential ECM modifier is the Matrix-Metalloprotease 2 (MMP2) which was shown to be expressed in the peripodial layer of the wing (Sui et al., 2012) and eye imaginal disc (Diwanji and Bergmann, 2020). Matrix Metalloproteinases are well known for their role in matrix degradation (Diaz-de-la-Loza et al., 2018; Sui et al., 2018, 2012). Consistently, overexpression of MMP2 in the wing disc results in a loss of the ECM shell, disc flattening and epithelial relaxation (Ma et al., 2017).

We stained wing discs for MMP2 and confirmed that MMP2 levels are higher in the PPE layer compared to the DP layer throughout disc growth (Fig.7A and Fig.S7A). We therefore hypothesized that peripodial MMP2 is required for planar growth of the top ECM_PPE_.

**Figure 7.**
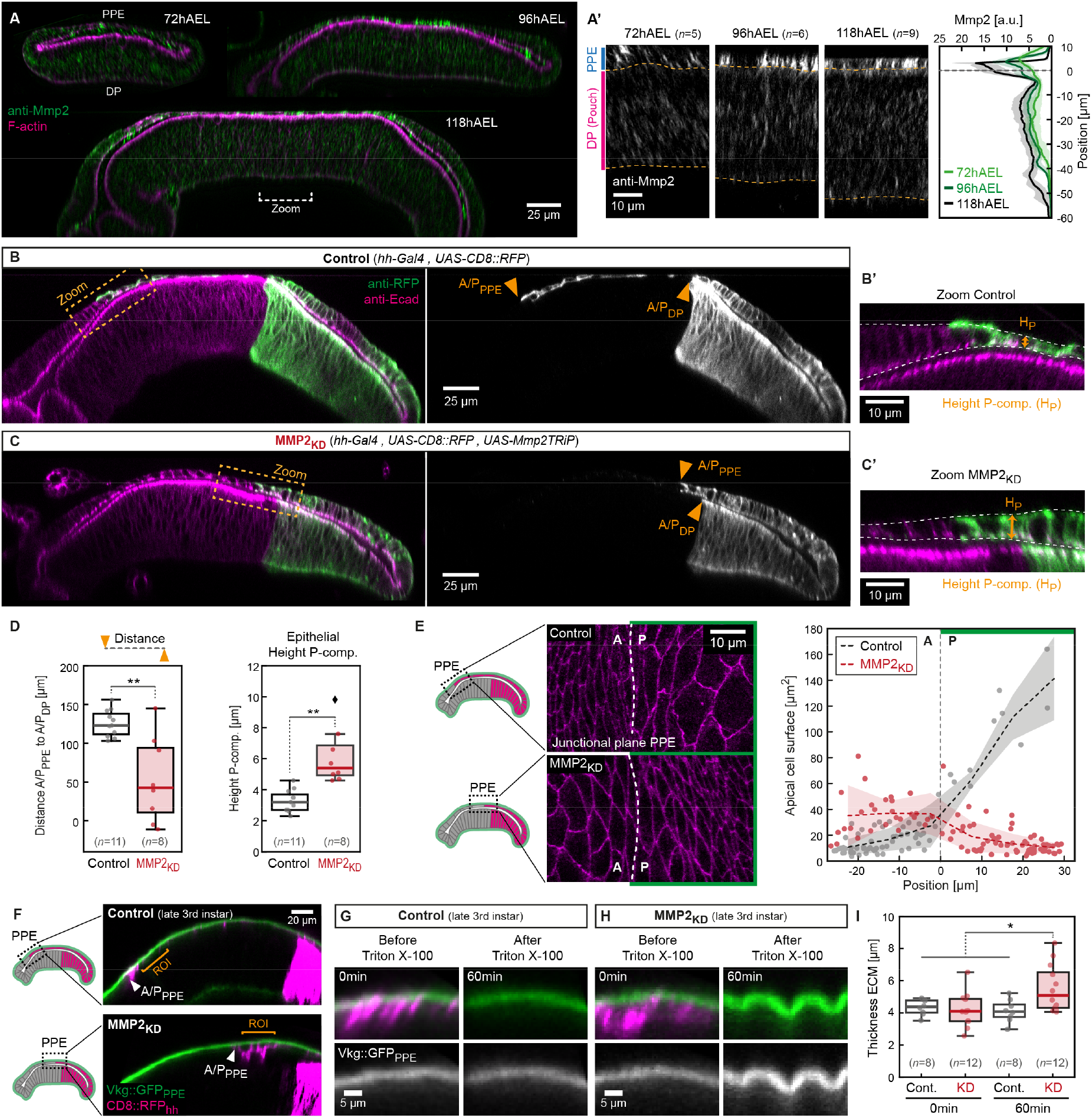
MMP2 modulates peripodial ECM growth anisotropy. (**A**) Section views of wing discs stained for MMP2 (green) at indicated time points and magnifications of the pouch region (A’, dashed lines mark DP). *right*: Quantification of average MMP2 fluorescence intensity distribution (dashed line indicates DP apical surface). (**B**) Section view of control wing discs expressing RFP (green) in the posterior compartment. A/P compartment boundaries are indicated in the PPE (A/P_PPE_) and in the DP (A/P_DP_) by arrowheads. (B’) Magnification of the A/P boundary region, indicated in (B). (**C**) MMP2_KD_ wing disc, the P-compartment in which MMP2 in knocked-down is marked by RFP (green). The A/P boundary region is magnified in (C’). (**D**) *left*: Quantification of the distance between the A/P boundary in the PPE versus the DP layer. *right*: Quantification of epithelial height (H_P_) as indicated in (B’+C’). (**E**) *left*: Representative magnifications of the peripodial A/P boundary region in x/y-view (junctional plane) in control (*top*) and MMP2_KD_ (*bottom). right*: Quantification of apical cell surface in the images shown left (error band indicates standard deviation). (**F**) Section of representative control (*top*) and MMP2_KD_ (*bottom*) wing discs expressing Vkg::GFP (green). The ECM_PPE_ was analyzed posterior of the peripodial A/P boundary (orange bracket, ROI). (**G**) Representative posterior section of a control peripodial ECM before (*left*) and after (*right*) decellularization. (**H**) Section of a MMP2_KD_ ECM before (*left*) and after (*right*) decellularization. (**I**) Quantification of peripodial ECM thickness of indicated conditions (assessed at ROI).

In order to investigate MMP2 function we knock-down (KD) MMP2 in the posterior compartment of the wing disc (referred to as MMP2_KD_, see Fig.S7B-C). MMP2_KD_ results in a posterior shift of the peripodial antero-posterior compartment boundary (A/P_PPE_) relative to the A/P boundary in the DP layer (A/P_DP_, see arrowheads in Fig.7B-C and Fig.7D, *left*). This shift coincides with increased epithelial thickness in posterior PPE cells where MMP2 is knocked-down (Fig.7D, *right*) and reduced apical cell surface area (Fig.7E). These results indicate that in MMP2_KD_ discs the planar expansion and flattening of PPE cells is inhibited and that PPE cells have a more compacted, cuboidal shape.

Next, we asked if reduced PPE expansion in MMP2_KD_ discs was due to changes in ECM growth anisotropy. Indeed, if growth of the ECM_PPE_ is no longer planar, the underlying PPE tissue layer is expected to be compressed in the plane and to thicken, as observed. As previously, we used decellularization to compare the relaxed ECM configuration in MMP2_KD_ and control discs. We decellularized late 3rd instar live explants (*ex vivo*) to directly follow changes in shape and intensity of the Vkg::GFP labeled ECM layer, focusing on the ECM_PPE_ posterior to the peripodal A/P_PPE_ boundary (see orange bracket in Fig.7F). Consistent with previous findings, decellularization of control discs did not result in changes in thickness (Fig.7G+I) or Vkg::GFP intensity (Fig.S7F). In contrast, in MMP2_KD_ discs decellularization resulted in a significant increase in ECM_PPE_ thickness (Fig.7H-I) and in Vkg::GFP_PPE_ intensity (Fig.S7D-F). Furthermore, the relaxed ECM thickness is significantly increased in MMP2_KD_ compared to controls (Fig.7I). Therefore, a loss of MMP2 modifies peripodial ECM_PPE_ growth anisotropy from planar to non-planar, 3D growth. In summary, we have shown that a spatial bias in MMP2 expression (high in the PPE and low in the DP) is required for planar growth of the top ECM_PPE_.

## Discussion

We investigated how growth of the wing imaginal disc affects its developing shape, namely tissue thickening and bending in a dome. Tissue doming emerges as the wing disc grows. Such doming is associated with the build-up of elastic stress as we report (Fig. 4 and 6). When this elastic energy is later relaxed by lysis of the PPE layer during pupariation (Aldaz et al., 2013; Pastor-Pareja et al., 2004), the wing imaginal disc everts and gives rise to the wing blade which further extends and flattens. Thus, we propose that tissue doming pre-stresses the wing disc for future metamorphosis. Our work sheds light on the mechanisms giving rise to tissue doming in response to growth.

Most studies so far have considered growth using indirect measurements of either cell proliferation or increase in cell number in clones over given time intervals (Mao et al., 2013; Pan et al., 2016). However, growth is intrinsically three-dimensional and requires a time dependent measurement of cell and tissue volume. Here, we quantified growth at the scale of few cells and the whole tissue in 3D over time and we provide evidence that growth is homogeneous within and between the epithelial layers of the wing disc. While spatial differences in growth rate of elastic material are well known to induce morphogenesis such as gut looping (Savin et al., 2011), villi formation in the chick (Shyer et al., 2013), gyrification in the cortex (Tallinen et al., 2016), this appears to play a limited role in the developing wing disc. One of our main findings is that the ECM, which forms an envelope around the disc, is not a passive boundary but an active, i.e. growing material. We have studied the relative volume growth as well as the anisotropy of growth of both the tissue and ECM layers of the disc. To disentangle the respective contributions of elastic deformation and growth, we used chemical methods to relax at different times the elastic constraint imposed by the ECM on the tissue (collagenase), and by the tissue on the ECM (TritonX-100). Strikingly, the two tissue layers grow in-plane, and the top ECM_PPE_ layer likewise, indicating compatible growth between the three top layers. In contrast, the bottom ECM_DP_ layer grows in 3D and its thickness increases. Therefore, control over the growth anisotropy of the bottom ECM layer is key to explain morphogenesis of the wing disc. However, epithelia can use active processes (e.g. adhesion and contractility) to actively control their height (Widmann and Dahmann, 2009). The fact that the stress-free configuration of the DP layer does not increase much in thickness over ~3 days argues that tissue thickening predominantly reflects elastic constraints of tissue growth imposed by the ECM shell rather than active regulation of cell surface tension by adhesion and cortical contractility.

We show that a nearly incompressible neo-Hookean model, in which pre-stress is added through differential growth, can well recapitulate 3rd instar wing disc development. Our modelling choice was guided by experimental results and relative simplicity. This is a valuable starting point to add more detail to the wing disc model, for instance by modelling explicitly all four tissue layers, modelling the hinge region, the ellipsoidal symmetry of the dome, and viscoelasticity. Furthermore, alternative stress-strain relationships than the neo-Hookean could be used to model, for instance, the strain-stiffening behavior observed in polymer networks like the ECM (Han et al., 2018).

What controls growth anisotropy of the ECM and what differs between the top and bottom ECM layers? Tissue-specific ECM biogenesis is complex and only starts to be revealed (Pastor-Pareja, 2020). The wing disc ECM consists of two interconnected networks of Laminin and Collagen IV in addition to other components. In *Drosophila* larvae the fat body was shown to be the main sources for most ECM components, such as Collagen IV (Dai et al., 2018; Pastor-Pareja and Xu, 2011). Therefore, the wing disc receives most ECM components via the hemolymph and their availability should not differ between the DP and the PPE. However, the disc tissue produces some portion of its Laminins (Dai et al., 2018; Urbano et al., 2009), suggesting that local production of ECM components might control differences in ECM structure. Here, we provide evidence that local production of MMP2 in peripodial cells is required for planar growth of the top ECM layer. We suggest that MMP2 modulates ECM turnover, possibly by digesting Collagen IV, thereby increasing the viscous dissipation of elastic energy due to growth of the underlying tissues. If the dissipation time scale of the top ECM_PPE_ layer is short compared to the growth time scale (cell doubling time ~8h), then the growing ECM layer is expected to flow and expand in-plane with the tissue and to remain in a stress-free configuration. However, if MMP2 levels are low, such as in the DP, the ECM crosslinking is expected to be comparatively higher, resulting in reduced viscous dissipation as the DP tissue grows such that the ECM grows in 3D. Therefore, the switch between planar versus thickness growth could be partially controlled by ECM turnover dynamics affecting ECM viscosity favoring either planar integration (high turn-over) or thickness growth (low turnover). Future studies will address this important problem.

In this context, the relative stiffness of the ECM_PPE_ and ECM_PP_ layer is not an essential mechanical parameter since the top PPE layer is in a stress-free, rest configuration. The most important feature is the differential growth anisotropy between the two ECM layers.

The ECM consists of a wide variety of molecules that influence their physical properties and response to stress (Dolega et al., 2021). Recently, the ECM component Hyaluronan was shown to induce osmotic ECM swelling during optic vesicle morphogenesis in zebrafish (Munjal et al., 2021) and in chick presomitic mesoderm (PSM) elongation (Michaut et al., 2022). Swelling results in ECM expansion deforming the overlying epithelial layer or expanding the PSM. Therefore, not only changes in remodeling rates, but also differences in molecular composition determine the material properties and response of ECM during development.

In this context, our study adds to the emerging notion that the ECM is a complex material whose growth is actively regulated and required for morphogenesis. As such it forms a geometric template that guides morphogenesis via a regulation of cell polarity and tissue mechanics. Our work underscores the fact that the elastic ECM enveloping the wing imaginal disc is a form of “active” shell whose growth affects the tissue layers as morphogenesis proceeds. While this is well documented in plants, fungi and bacteria where the cell wall is synthesized and modified to guide osmotic growth of cells and tissues, this is not much appreciated in animal morphogenesis. As such, through the regulation of the ECM by e.g. MMP2 and possibly other enzymes, organs have the potential to modify the geometric and mechanical information contained in the ECM shell. It will be important to further explore how such feedbacks operate during morphogenesis.

## Acknowledgements

We would like to thank all the members of the Lecuit team for stimulating discussions. We also thank Larry Taber (Washington University in St. Louis) for instructions and sample codes in the implementation of growth in a finite element framework. We are grateful to B. Aigouy (IBDM, France) for help with Tissue Analyzer,Benoit Dehapiot (CENTURI, France) for help with Python and to Stephane Noselli (IBV, France), Frank Schnorrer (IBDM, France), Kendal Broadie (Vanderbilt, USA), the Bloomington Stock Center and the Developmental Studies Hybridoma Bank (University of Iowa, USA) for flies and reagents. The IBDM imaging platform and the France-BioImaging infrastructure supported by the Agence Nationale de la Recherche (ANR-10-INSB-04-01; call “Investissements d’Avenir”) provided support. This work was supported by the Ligue Nationale Contre le Cancer (Equipe Labellisée 2018).

S.H. was supported by an EMBO long-term fellowship (ALTF 217-2017), the College de France (Paris, France). S.H and A.E. were supported by postdoctoral fellowships from the Turing Center for Living Systems (CENTURI), funded by France 2030, the French Government program managed by the French National Research Agency (ANR-16-CONV-0001) and from Excellence Initiative of Aix-Marseille University - A*MIDEX ».

## Author contribution

S.H., A.E., G.Z. and T.L. conceived and designed the study. S.H. performed the experiments and quantified the data. A.E. designed the computational model and performed the simulations. All authors discussed the data. S.H., A.E. and T.L. wrote the manuscript. All authors read and approved the manuscript.

## Material and Correspondence

A supporting text describing the modeling and fitting procedures is available <u>here</u>.

Correspondence and material requests to Thomas Lecuit -

## STAR Methods

### 1. Fly strains

The following fly lines were used: *y^1^,w^1118^, hs-Flp; act>Stop>Gal4, UAS-EGFP* (AyGAL4, originating from Bloomington stock 64231); *UAS-Histone::mRFP* (Wirtz-Peitz et al., 2008), *Vkg^G454^::GFP* (Morin et al., 2001) (both from F. Schnorrer), *vgQE-dsRed* (Zecca and Struhl, 2007)*, sqh-Sqh::mCherry* (Bailles et al., 2019) (insertion site 53B2), *endo-Ecad::GFP* (Huang et al., 2009). The following lines were obtained from the Bloomington stock center: *AGiR-Gal4* (#6773), UAS-PI3K_DN_ (#25918), UAS-CD8::mRFP (#27398), UAS-CD8::RFP (#27392), UAS-Mmp2 TRiP (#61309), Mi{MIC} insertion in Mmp2 (#60512). *hh::Gal4* is described on FlyBase (www.flybase.org).

### 2. Genotypes by figure

Figure 1: *y^1^,w^1118^*
Figure 2: (**C+D**) *hs-Flp; tub>Stop>Gal4, UAS-EGFP / UAS-Histone::mRFP*
Figure 3: (**C**) *hs-Flp; tub>Stop>Gal4, UAS-EGFP / UAS-Histone::mRFP*
Figure 4: *Vkg^G454^::GFP / endo-Ecad::GFP, sqh-Sqh::mCherry*
Figure 5: *Vkg^G454^::GFP* / +
Figure 6: (**D**) *y^1^,w^1118^*
Figure 7: (**A**) *y^1^,w^1118^*, (**B-E**) w; UAS-Mmp2TRiP / +; hh-Gal4 / UAS-CD8::RFP, (**F-I**) w; UAS-Mmp2TRiP / Vkg::GFP; hh-Gal4 / UAS-CD8::RFP

Figure S1: *y^1^,w^1118^*
Figure S2: (**A-C**) *hs-Flp; tub>Stop>Gal4, UAS-EGFP / UAS-Histone::mRFP*, (**D-E**) *y^1^,w^1118^*
Figure S3: (**A**) *y^1^,w^1118^*, (**C**) *UAS-CD8::mRFP / +; AGiR-Gal4* / +, (**D**) *UAS-CD8::mRFP / UAS-PI3K_DN_; AGiR-Gal4* / +
Figure S4: (**A+C**) *Vkg^G454^::GFP / endo-Ecad::GFP, sqh-Sqh::mCherry*
Figure S5: *Vkg^G454^::GFP* / +
Figure S6: *Vkg^G454^::GFP* / +
Figure S7: (**A**) Mi{MIC} insertion in Mmp2 (#60512), (**B-C**) w; UAS-Mmp2TRiP / +; hh-Gal4 / UAS-CD8::RFP, (**D**) w; UAS-Mmp2TRiP / Vkg::GFP; hh-Gal4 / UAS-CD8::RFP

### 3. Antibodies

Primary antibodies used were mouse-anti-Wingless (4D4-s; 1:120; DSHB, University of Iowa); mouse-anti-Patched (Apa1-s; 1:40; DSHB, University of Iowa); rat-anti-DE-cadherin (DCAD2 concentrate; 1:200; DSHB, University of Iowa); rabbit-anti-GFP (1:1000, Abcam ab6556); rabbit-anti-Phospho-Histone H3 (PHH3, 1:1000, Cell Signaling #9701); rabbit-anti-Mmp2 (1:500, from K. Broadie (Dear et al., 2015)).

Tissue outlines were marked by Alexa Fluor 660 Phalloidin (1:50, A 22285, Sigma Aldrich) which was added together with the other secondary antibodies. Secondary antibodies from the AlexaFluor series (Sigma Aldrich) were used at 1:500 dilution. Discs were blocked in 2% normal donkey serum (017-000-121, Jackson ImmunoResearch).

### 4. Sample collection, immunostaining and imaging

For staged samples, embryos were collected for 2h intervals as described before (Harmansa et al., 2015) and allowed to develop at 25°C until the desired developmental stage (MMP2 knock-down experiments in Fig.7B-I were performed at 29°C due to increased efficiency of knock-down). Wing discs were isolated at defined time intervals after egg laying (hAEL). For 72hAEL and older time points only male larvae were included (positive selection by the transparent genitalia disc well visible in the posterior half of male larvae); 65hAEL data contains male and female larvae since at this time point the genitalia disc is not yet clearly visible.

All larvae of one experiment were dissected, processed and imaged in parallel, using identical solutions in order to reduce experimental variations. Immunostaining of imaginal discs was performed as described previously (Harmansa et al., 2015). Discs were mounted in Vectashield Plus (H-1900, Vector Laboratories) using double sided tape as spacers (TESA 05338) to maintain tissue morphology and avoid squishing of the sample.

All fixed samples were imaged on a Leica SP8 confocal microscope using a 40x or a 63x/1.4 NA oil-immersion objective. All image stacks of one experiment were acquired in the same session using identical imaging settings. Imaging conditions were chosen to be well within the dynamic range of the fluorescent signal obtained. For optical cross sections of wing discs stacks with high resolution along the z-axis were acquired (typically 0.33μm spacing between slices).

### 5. Image processing

Image data was processed and quantified using Fiji/ImageJ software (National Institute of Health). Further data processing was performed in Python. Individual procedures are described in detail in the following:

#### 5.1 Epithelial thickness and bending quantifications

Image stacks of high resolution along the z-axis of discs stained for Wingless (Wg) and Patched (Ptc) were acquired (typical spacing between slices are 0.33μm). Stacks were sliced using the ‘Reslice [/]’ function in Fiji to obtain optical cross sections parallel to the dorsal/ventral boundary with a slight dorsal offset. Thickness of the disc proper layer and the overlaying peripodial layer were measured at the position of the cross marked by the horizontal Wg and the vertical Ptc expression using the ‘Straight line’ tool.

In order to visualize the average basal shape of the disc proper epithelium, the basal outline of the disc proper epithelium was marked using the ‘Kappa - Curvature Analysis’ plugin in Fiji. Kappa allows the export of a spline-fit the basal surface outline. Basal outlines were registered along the x-axis, defining 0 as the position of the A/P boundary (marked by Ptc). Registered profiles were subsequently fitted in Python by a B-spline using a custom script. In plots, the average basal outline is depicted by a solid red line and profiles of individual discs in gray.

#### 5.2 - 3D segmentation and volume qualifications (Fig.S1)

In order to assess volume growth at the tissue level we have focused on the wing pouch in the DP epithelium. We have used antibody stainings against Wingless (Wg) and Patched (Ptc) to mark the pouch by the inner ring of Wg expression. We have followed two approaches to obtain volume information of the wing pouch:

1. First we have performed proper 3D segmentation of volumetric image stacks of staged wing discs (72 and 118hAEL) stained for Wg/Ptc (see Fig.S1, E-G). The Wg/Ptc staining results in sufficient labeling of epithelial outlines. We segmented the epithelial volume (Wg/Ptc signal) in Ilastik (Berg et al., 2019) using the ‘pixel classification’ and obtaining a binary mask of the segmented epithelial signal. The binary mask was manually corrected for errors in Fiji and finally restricted to the volume surrounded by the inner Wg ring. Volume values for the segmented wing pouch were obtained from the binary mask using the ‘Histogram’ function and by multiplying the obtained number of wing pouch pixels with the voxel volume. Indeed, this procedure is work intensive since it requires a significant amount of manual correction in Fiji.
2. We therefore have tried to approximate the wing pouch volume by simply multiplying pouch epithelial thickness with the area of the inner Wg ring (see Fig.S1, A-D). Epithelial thickness was measured close to the intersection of the Wg/Ptc cross. Wg ring area was measured in maximum projections using the ‘Polygon selection’ tool in Fiji. In order to correct for the increased area due to tissue doming after 80hAEL we have approximated the Wg ring area as spherical cap with cap height *h* (which was obtained from the average basal outlines in Fig.1B’). Please see Fig.S1B+C for details on this correction. Indeed, this approximation yields values very close to the 3D-segmented ‘true’ values for both the flat 72hAEL and the bend 118hAEL time points (see Fig.1G). We therefore used the less work intense approximation approach to quantify wing pouch volume in Fig.1D’.

Analogous to procedure (1) we segmented the volume of the peripodial epithelium overlying the wing pouch (see Fig.S1H-J). Due to a lack of landmarks in the peripodial layer, we have decided to quantify the volume of the peripodial tissue that covers the inner Wg ring. Hence peripodial volume values plotted in Fig.S1J correspond to the peripodial volume that covers the wing pouch tissue.

#### 5.3 Quantification of average cell volume (Fig.S2)

In order to obtain spatial and temporal information of average cell volume we imaged clones labeled by cytosolic EGFP and Histone::RFP (as described before) with high z-resolution. The obtained EGFP signal was subsequently segmented using Ilastik (Berg et al., 2019) (‘pixel classification’ providing a binary output image) and 3D volume per clone was obtained using the ‘3D manager’ of the ‘3D imaging suit’ in Fiji. Obtained clonal volumes were divided by the number of cells per clone (obtained from the Histone::mRFP labeling) to obtain the average cell volume. Data from multiple discs were spatially averaged using the Wg/Ptc landmarks.

#### 5.4 Quantification of clonal proliferation rates (Fig.2C)

Clones were induced by heat-shock induced cassette recombination that resulted in clonal expression of EGFP and Histone::RFP (*hs-Flp; act>Stop>Gal4, UAS-EGFP / UAS-Histone::RFP*). As a general rule, wing discs were isolated 24h after clone induction, therefore staged larvae were heat shocked (HS) at 37°C at defined time points and dissected 24h later (hence HS at 48hAEL for 72hAEL samples). HS length was shorter for early time points (4min for 72hAEL sample) and longer for older samples (7 min for 116hAEL) to obtain discs with sparse clonal density in order to reduce clone fusion and associated mistakes in estimating clonal proliferation rates. Isolated discs containing clones were stained for GFP, RFP and Wg/Ptc (landmarks) and imaged using a Leica SP8 confocal microscope at 63x magnification. Nuclei per clone were counted using the Fiji ‘3D viewer’ and the ‘orthogonal views’ tools in order to correctly assess nuclear numbers in 3D. Given that each clone originates from a single founder cell, the cell number *n* after 24h of clone induction allows us to calculate the number of proliferation events that have taken place within these 24h using log2(*n*). Average spatial proliferation maps were created by arranging the data according to the landmarks provided by the Wg/Ptc staining.

#### 5.5 Quantification of proliferation rates via PHH3 (related to Fig.S3A)

Wing discs (*y^1^,w^1118^*) were isolated at defined time points of development and stained for Phospho-Histone H3 (PHH3, marker for mitosis), E-cadherin (cell outlines), and Wg/Ptc (serving as landmarks for registration of multiple discs). Image stacks were obtained on a Leica SP8 microscope using a 63x objective. Apical surface projections of the disc proper surface using the E-Cad signal were obtained using a custom made Fiji plugin based on the ‘Stack Focuser’ plugin. Subsequently, cell outlines were segmented using the ‘Tissue Analyzer’ plugin (Aigouy et al., 2016). The landmarks provided by the Wg/Ptc cross and ring were used to register multiple discs and to obtain average spatial distributions of cell area and cell density for each time class. As published previously (Mao et al., 2013), we observe that the cell area becomes non-uniform around 80hAEL with smaller cells in the center compared to the periphery of the wing pouch (not shown). Average proliferation profiles were created based on the PHH3 signal. Given that cell density is not uniform in space, we subsequently binned the cell density and proliferation density profiles in rectangular regions of 8μm edge length. Proliferation density profiles were normalized per bin to obtain the average proliferation rate per cell in a spatial manner. In order to investigate spatial non-uniformities in proliferation between the center and the periphery of the disc we divided the wing disc in 4 elliptic rings within the region marked by the inner Wg ring. While the central 3 regions correspond to wing pouch tissue, the outermost region corresponds to hinge tissue (see Fig.S3A, *right*). Importantly, independent of the method chosen for data quantification, we never obtained higher proliferation values in the center versus the periphery. In contrast, correct normalization of cell proliferation by cell density shows a tendency of decreased central proliferation after 80hAEL.

#### 5.6 Extraction of Vkg::GFP concentration profiles and ECM thickness

We have used the Vkg::GFP signal to assess changes in ECM structure and thickness. In order to quantify absolute changes in Vkg::GFP levels and distribution we have acquired image stacks with high z-resolution of either the top ECM_PPE_ or the bottom ECM_DP_. Importantly, we imaged either the top or bottom ECM depending on the orientation of the wing disc and which ECM (DP or PPE) was closer to the objective after mounting. Optical cross-sections of image stacks close to the Wg/Ptc intersection were obtained using the ‘reslice’ function in Fiji. From average projections of five consecutive slices we extracted the Vkg::GFP profiles using the ‘straight line tool’ (3.6μm width) and the ‘plot profile’ function in Fiji at three random positions (in order to average out small local differences). These three profiles were aligned by the position of their peak intensity and averaged. Finally, average profiles from multiple wing discs were averaged in order to obtain representative Vkg::GFP profiles for the top ECM_PPE_ and the bottom ECM_DP_ for the different developmental time points (see Fig.S5, B’ and D’). Profiles were plotted in Python using the Seaborn package (‘lineplot’ command). In the average profiles the dashed line indicated the average fluorescent intensity and the error bands the standard deviation.

In order to quantify changes in ECM thickness we chose intensity thresholds at values that capture most of the observed Vkg::GFP fluorescence (see Fig.S7, B’ and D’, threshold values = 10a.u.). We measured the width of the Vkg::GFP profiles at the given threshold in order to compare changes in thickness between different time classes and experimental treatments (see Fig.S5, B’’ and D’’). Analogously, we extracted profiles in *ex vivo* cultured discs upon decellularization (See Fig.S6C+D). The only difference were the chosen intensity threshold values (10a.u. for the ECM_DP_ and 15a.u. for the ECM_PPE_) that differed due to the different imaging conditions of the live sample.

### 6. Details on *ex vivo* culture and imaging (Fig.2+3)

The procedure of long-term imaging of disc explants was based on the protocol published by (Dye et al., 2017) with minor modifications. We slightly modified the composition of the culture medium by adding adenosine deaminase (ADA, 8.3 ng/ml final concentration, Roche 10102105001) as proposed by recent findings of (Strassburger et al., 2017). In our hands, the addition of ADA in particular improved the long term culture of young disc explants. In contrast, the addition of juvenile hormone (Methoprene) as proposed by Strassburger *et.al*. has not proven beneficial in our setting and we did not use it in our culture medium.

As described previously (Dye et al., 2017), 72hAEL old larvae were dissected in culture medium and explants were immobilized between a round coverslip and a porous filter membrane (Whatman cyclopore polycarbonate membranes; Sigma, WHA70602513) using double sided tape as spacers (~50μm thickness, 3M Scotch ATG 904 Clear Transfer Tape, No. 909-3799 from RS Components). The coverslip containing the mounted explants was inserted in an Attofluor chamber (A7816, ThermoFisher) and filled with 1ml of culture medium. Explants were imaged on a Nikon Roper spinning disc Eclipse Ti inverted microscope using a 40X-1.25 N.A. water-immersion objective at 22°C. Image stacks of 1μm z-spacing were acquired in 10 min intervals for up to 12 hours.

#### 6.1 Quantification of volume growth rates

We staged larvae of the genotype *hs-Flp; act>Stop>Gal4, UAS-EGFP* to 72hAEL. Clonal expression of a cytosolic GFP was induced by heat shock (at 37°C for 4min) 1h before dissection (60hAEL). Wing discs of 72hAEL old animals were isolated, cultured and imaged as described before. Individual clones from volumetric movies were cropped in Fiji, background was subtracted (using a ‘rolling ball’ radius of 50 pixel) and the volume marked by the cytosolic GFP signal segmented using a ‘pixel classification’ in Ilastik (Berg et al., 2019). Clonal volumes were assessed for each hour of the movie by averaging three consecutive timepoints and the hourly growth rate was calculated (see plots in Fig.2D’). For each clone we calculated an average growth rate for the full span of the movie (up to 10h). In order to compare spatial differences in growth rates in the plane of the DP epithelium we grouped central clones, defined as clones within an ellipse covering the central 60% of the wing discs (see scheme Fig.2E) and compared their growth rate to peripheral ones. Analogously, we segmented and analyzed growth rates in the peripodial epithelium.

### 7. Correlating relative tissue with ECM growth (Fig.S5)

In order to gain understanding of the volumetric increase of the epithelial compared to the ECM layers we plotted the relative volume increase of the tissue versus the relative increase in the ECM layer. For the DP, volume was quantified in Fig.1D’, and normalized by the average value either at 65hAE or at 72hAEl. In the time interval between 65-118hAEL the DP volume increases by ~65.8-fold and between 72-118hAEL by ~19.9-fold. For the bottom ECM_DP_ layer we first estimated the volume by multiplication of the known area of the inner Wg ring (see Fig.S1) with the known ECM_DP_ thickness (Fig.S5C’’). Estimated ECM volume increases ~16.2-fold from 72 to 118hAEL.

In addition, we quantified Viking::GFP integrated intensities of the signal lining the basal side of the wing pouch (area within the inner Wg ring). For this we created a temporal data set of disc of the genotype Vkg::GFP / +. Processing (fixation, immunostaining and mounting) was done under identical conditions using identical solutions. Subsequently, the mounted discs were imaged in one session using identical settings to allow direct comparison of fluorescent Vkg::GFP intensities. Only discs with their basal side of the DP facing towards the objective were imaged and included in the quantifications. For processing (in Fiji), the Vkg::GFP fluorescent signal was restricted to the volume lining the inner Wg ring and after background subtraction (rolling ball with radius = 50) the integrated Vkg::GFP intensity was calculated using the ‘histogram’ function. We found that the integrated fluorescence intensity of the Vkg::GFP marked ECM_DP_ increases by ~56.6-fold between 65-118hAEL. Consistently, both approaches, ECM volume estimation and integrated intensity measurements, suggest that the volumetric growth of the ECM is reduced compared to the overlaying DP tissue.

We performed the same analysis for the peripodial layer. Given the lack of landmarks in the PPE layer, we decided to include the peripodial volume that overlays the inner Wg ring (corresponding to the wing pouch) in our quantifications. The peripodial volume overlaying the Wg ring increased by ~8.2-fold between 72-118hAEL (see Fig.S1J). In contrast, the ECM_PPE_ volume, estimated by multiplying the Wg ring area with the ECM_PPE_ thickness (see Fig.5B’’), increased by ~7.9-fold on average, a value very similar to the overlaying PPE cell layer.

### 8. Acute ECM modifications and decellularization

#### 8.1 Acute ECM digestion using Collagenase (Fig.4 and S4)

Larvae of the genotype *Vkg^G454^::GFP / endo-Ecad::GFP, sqh-Sqh::mCherry* were staged as described above. Larvae were dissected in PBS and transferred to Eppendorf tubes containing PBS on a 37°C heat block. In order to inhibit Myosin II activity the ROCK inhibitor Y-27632 dihydrochloride (Sigma-Aldrich, Y0503) was added to a final concentration of 2mM and incubated for 2 minutes. Subsequently, the extracellular matrix was digested by addition of Collagenase (Sigma-Aldrich, C0130) at a final concentration of 1mg/ml and incubated for 1 minute. After 1 minute of Collagenase treatment, discs were fixed by direct addition of fixative (4% PFA in PBS) to maintain and conserve disc morphology after ECM digestion. Discs were fixed for 20 minutes at RT on a rocker and subsequently processed for immunostaining as described above.

#### 8.2 Decellularization of wing discs (Fig.5 and S6)

Here, we have adopted and used a chemical decellularization method to free the wing disc extracellular matrix from the load exerted by the epithelial cell layers. While classical decellularization protocols often use strong detergents like e.g. SDS, wing disc cells are soft and increased concentrations of Triton X-100 are sufficient to permeabilize and degrade cells. *Vkg^454^::GFP* larvae were staged to 72, 96 and 118hAEL to cover the whole 3^rd^ instar development. Larvae were dissected in PBS. For decellularization, dissected and inverted larvae were incubated in PBS + 3% Triton X-100 for 15 minutes before fixation, while control discs were incubated in PBS. Shorter exposure to Triton X-100 did not result in sufficient separation of the ECM layer from disc proper cells; more than 15 minutes resulted in a loss of the cell layer and hence a loss of the required landmarks to identify peripodial versus disc proper ECM layers. After fixation, discs were processed for immunofluorescence as described before.

All discs were mounted on the same microscopy slide and imaged under identical conditions. For each wing disc, depending on its orientation, the ECM closer to the objective was imaged (either peripodial or disc proper ECM). Hence, only image data acquired close to the objective was included in intensity quantifications to avoid inaccuracy due to loss of signal with increasing imaging depth.

Optical cross sections of the region around the intersection of the A/P D/V boundaries were obtained by slicing the image stack using the ‘reslice function’ in Fiji/ImageJ. Intensity profiles along the apical-basal axis of the Vkg::GFP signal were obtained using the ‘straight line tool’ (line width of 3.6μm). Multiple profiles were aligned according to their peak intensity, averaged and plotted in Python using the seaborn library (line plot function, error bands represent the standard deviation). Thickness changes of the ECM layer under load and upon relaxation were quantified as described in section 5.6.

#### 8.3 Circular bleaching and *ex vivo* decellularization (Fig.5D and Fig.7F-H)

Decellularization results in a loss of the epithelial layers and hence landmarks provided by the cell layers (like e.g. the Wg/Ptc ring and cross). In order to assess relaxation in the x/y-plane of the ECM we used a photobleaching approach to mark circular regions that could serve as traceable landmarks that are not lost during decellularization.

Vkg::GFP wing discs of 116hAEL were isolated in PBS and glued to the bottom of a petri dish using classical embryo glue. Embryo glue was applied shortly before mounting, briefly allowed to dry and then covered by a drop of PBS in which the discs were arranged. For experiments assessing the ECM_DP_, discs were glued with their PPE side to the bottom of the petri dish, their ECM_DP_ facing upwards. For assessing the ECM_PPE_ discs were mounted with inverse orientation.

Experiments were performed on an upright Nikon A1R MP+ multiphoton microscope. Mounted live discs were taken directly to the microscope and imaged from the top using a water immersion objective (40x/1.15NA). For excitation of GFP a tunable wavelength pulsed laser (Coherent) at 920 nm was used. Imaging settings were optimized to use minimal laser power. Circular regions of interest (ROI, usually 3 circles per disc) were bleached using elevated laser power and scanning the ROI for 30-times. Depending on the geometry of the ECM this was repeated for multiple positions along the z-axis to obtain a clear circle upon maximum projection of an image stack (1μm spacing). The circular bleaching procedure was restricted to a total of 40min (~5 discs per session) such that including the mounting time the total time of discs in PBS did not exceed one hour before decellularization. After marking circles on all discs, the petri dish was filled with PBS containing 3% Triton-X100 (PBST-3%). Discs were imaged before addition of Triton X-110 and subsequently in 10min intervals after exposure to Triton X-100 in order to follow ECM shape changes due to the loss of constraints induced by the cell layers. Obtained image stacks were subsequently oriented in Fiji (using the reslice and transformation functions) to ensure that circles are not tilted but are in-plane with the projection plane. Circular area was measured in Fiji using the ‘polygon selection tool’ in maximum projections. Relative area changes were processed in Excel and plotted in Python (Seaborn library).

In order to investigate ECM thickness changes upon decellularization, samples were prepared the same way, however, per disc only 2 circles were bleached and image stacks with high z-resolution were obtained (0.25μm spacing). 60min after exposure to Trition X-100 another set of high z-resolution image stacks were acquired under identical settings. Subsequently, ECM thickness profiles were processed as described in section 5.6.

### 9. Quantification of changes in peripodial cell architecture upon Mmp2KD (Fig.7B-E)

The peripodial layer is thinner and shows lower fluorescent signal (of e.g. Ecad or RFP) than the DP layer. Even after optimization of the Stack Focuser plugin we were not able to obtain satisfiable results for apical surface projections of the peripodial layer. We therefore manually create a mask for the junctional plane of the PPE layer. This was done in Fiji creating an additional channel (‘mask_PPE_’) in which the peripodial surface was marked using the pencil tool such that in the final mask_PPE_ stack pixels either had a value of 1 if they correspond to the peripodial apical surface or a value of 0 otherwise. Multiplying the mask stack with the Ecad stack (using the ‘image calculator’ function) allowed us to extract only peripodial Ecad signal which after maximum projection yielded the PPE junctional plane. Cell outlines in PPE projections were then segmented in the region of the A/P boundary using Tissue Analyzer (Aigouy et al., 2016).

Epithelial thickness was measured in cross-sections obtained by using the ‘reslice’ tool in Fiji. Thickness was measured using the ‘straight line’ tool (Fiji) 10μm posterior of the peripodial A/P boundary (as indicated in Fig.7B’ and C’). In the same cross-sections the distance between the peripodial and the disc proper A/P boundary was measured using the ‘segmented line’ tool.

### 10. Statistics and data representation

Given the experimental constraints we aimed to obtain a sample size large enough (*n* ≥ 5) to allow testing statistical significance by using a two-sided Student’s *t*-test (unequal variance, \**p*≤0.05, \*\**p*≤0.005, \*\*\**p*≤0.0005). The number of samples and *p*-values are either indicated in the figure or the respective legend. For each experiment *n*-numbers indicate biological replicates, meaning the number of biological specimens evaluated (e.g. the number of wing discs or clones). Plots were created in Python using the Seaborn library. In line plots the error bands indicate the standard deviation. In box plots the median is indicated by a central thick line while the interquartile range (containing 50% of the data points) is outlined by a box. Whiskers indicate the minimum and maximum data range, outliers are indicated by a black rhomb and were excluded from further processing.

## Supplementary information titles and legends

**Figure S1.**
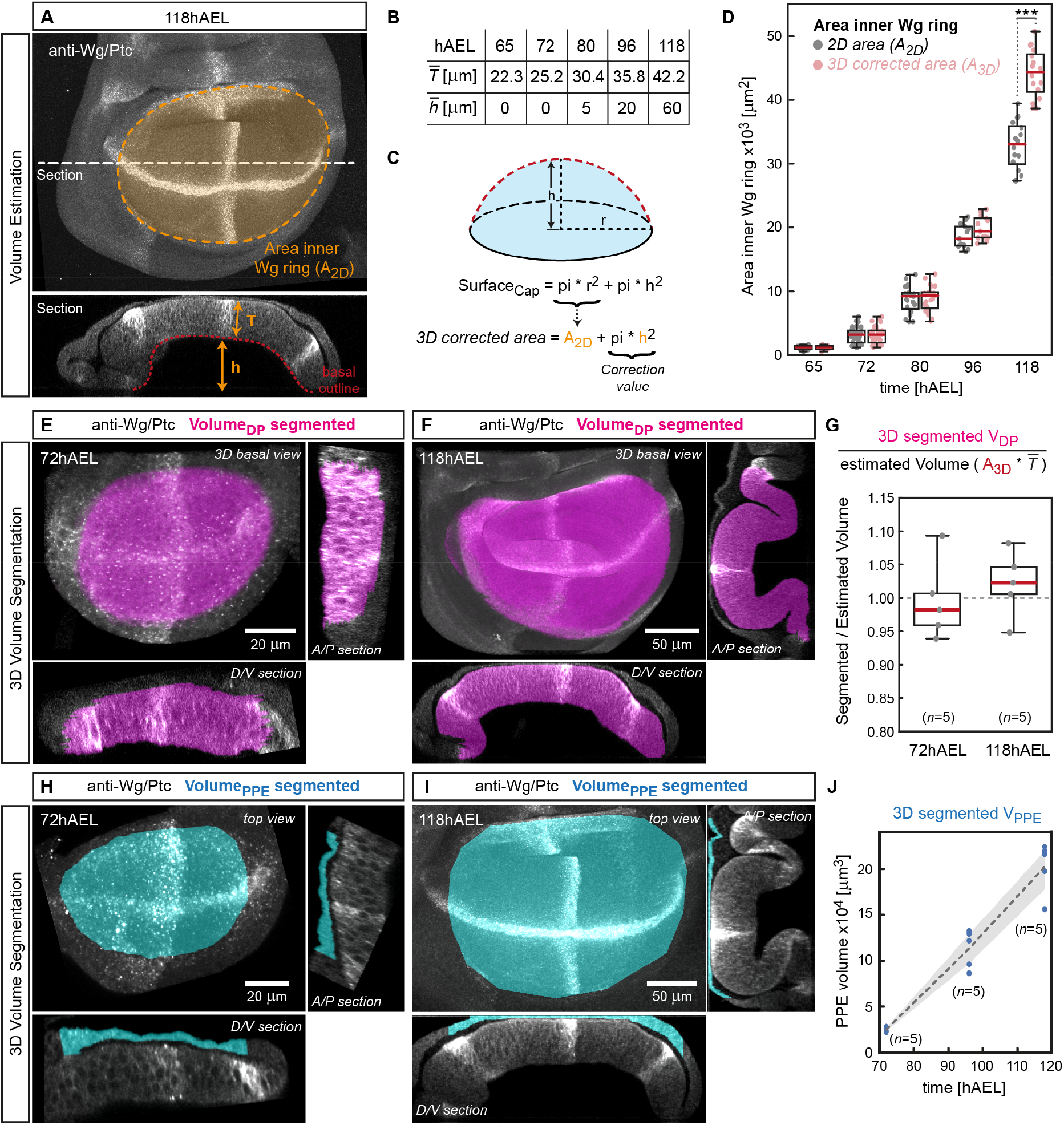
Measuring volume growth of the disc proper and peripodial epithelium. (**A**) Projection of a 118hAEL wing disc stained for Wg/Ptc. The wing pouch area can be estimated from the region outlined by the inner Wg ring (A_2D_, marked by orange dashed line). In the cross section view (bottom) we can measure the thickness *T* of the DP epithelium and the height *h* of the ‘dome’ that the disc forms at stages after 80hAEL. (**B**) Table of average values for DP thickness *T* and dome height *h* at indicated times. DP thickness has been measured as indicated close to the A/P-D/V intersection (see panel A bottom). Average height of the dome was obtained from the basal surface profiles in Fig.1B’. (**C**) In order to correct the measured projected 2D area (A_2D_) for the 3D effect of doming we have assumed that the basal surface of the disc after 80hAEL resembles a spherical cap. The surface of a spherical cap can be calculated knowing the base area (in our case A_2D_) and the height of the dome *h*. Therefore, the 2D measured area values can be corrected by adding pi\**h*^2^, a value that is fixed for a given developmental stage. (**D**) Comparison of wing pouch area measured in 2D (A_2D_, grey) and corrected for doming (A_3D_, red). In particular at 118hAEL this correction makes a significant difference. (**E+F**) In order to obtain precise values for the wing pouch volume we have segmented the volume of the DP tissue encircled by the inner Wg ring in 3D (see methods) at 72hAEL (**E**) and 118hAEL (**F**). Shown is a 3D volume view from the basal side of the DP with the segmented 3D volume marked in magenta (*top left*) and section views along the A/P boundary (*right*) and the D/V boundary (*bottom*). (**G**) 3D segmentation of the pouch volume as shown in E and F requires significant manual correction work. We therefore evaluated to which extent an estimate of DP volume (obtained by multiplying the 3D corrected projection area (A_3D_) with the thickness T of the DP epithelium) can match the actual DP volume values obtained by 3D segmentation. For this we segmented the DP volume of 5 discs at 72hAEL (flat tissue) and at 118hAEL (domed tissue) and compared the obtained values with the estimated (A_3D_\**T*) values. Indeed, the estimated values are very close to the actual volume values, with an average error of only ~4%. We therefore conclude that estimation of the Dp volume (A_3D_\**T*) yields precise values that are sufficient for investigating volumetric growth rates of the DP tissue. (**H-J**) Segmentation of the peripodial volume overlaying the wing pouch (using the inner Wg ring as landmark). Example discs at 72hAEL (H) and at 118hAEL (I) labeled for Wg/Ptc and the segmented PPE volume is marked in cyan. The volume increase between 72 to 118hAEL is plotted in (J) (*n*=5 for each time point, the error band represents the standard deviation).

**Figure S2.**
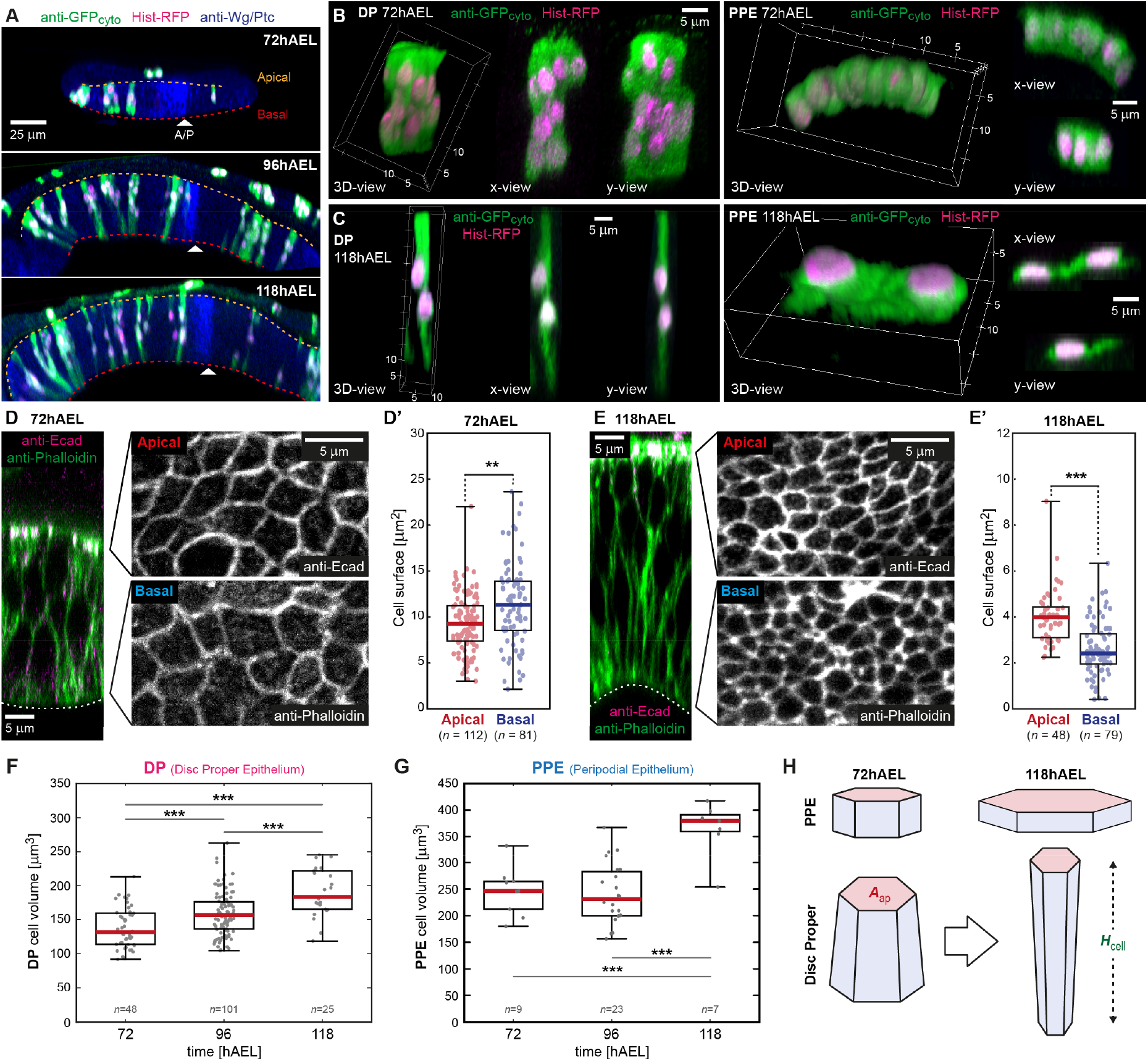
Cell shape changes associated with tissue doming. (**A**) Section view of discs at 72hAEL (top), 96hAEL (middle) and 118hAEL (bottom) expressing cytosolic GFP (*GFP_cyto_*, green) and *Histone::RFP (Hist::RFP*, magenta) in clones of cells. Clones were induced 24h before dissection, e.g. at 48hAEL for the 72hAEL sample. The apical and basal surface of the DP are marked by dotted lines (yellow and red, respectively). (**B**) Representative clones in the DP epithelium (left) and the PPE (right) at 72hAEL (24h after induction). The cell volume is marked by cytosolic GFP (*GFP_cyto_*, green) and nuclei are labeled by *Hist::RFP* (magenta). Each panel shows a 3D view of the clone (left) and section views in x- and y-direction (middle-right). (**C**) Same as in (B) but at the end of larval development at 118hAEL. (D) *left*: Section of a 72hAEL old wing disc close to the A/P-D/V intersection stained for E-cadherin (magenta) and Phalloidin (green). The basal surface is marked by a dashed line. *right*: Projections of the apical (top) and basal surface (bottom) to visualize changes in cell area. (**D’**) Quantification of apical and basal cell surface area at 72hAEL. At this stage the basal cell surface is slightly bigger than the apical one. (**E**) Section (left) and apical and basal projections (right) of an 118hAEL old wing disc as described in (A). (**E’**) Quantification of apical and basal cell surface at 118hAEL. At this stage the disc is doming upwards and the apical cell surface is bigger than the basal one. (**F**) Quantification of average cell volume in the disc proper at indicated time points (see methods). (**G**) Average cell volume in the PPE. (**H**) During wing disc growth cell shape in the DP and the PPE change drastically. While in the DP cells decrease their apical surface (A_ap_) and increase their height (H_cell_), in the PPE cells flatten and become more squamous.

**Figure S3.**
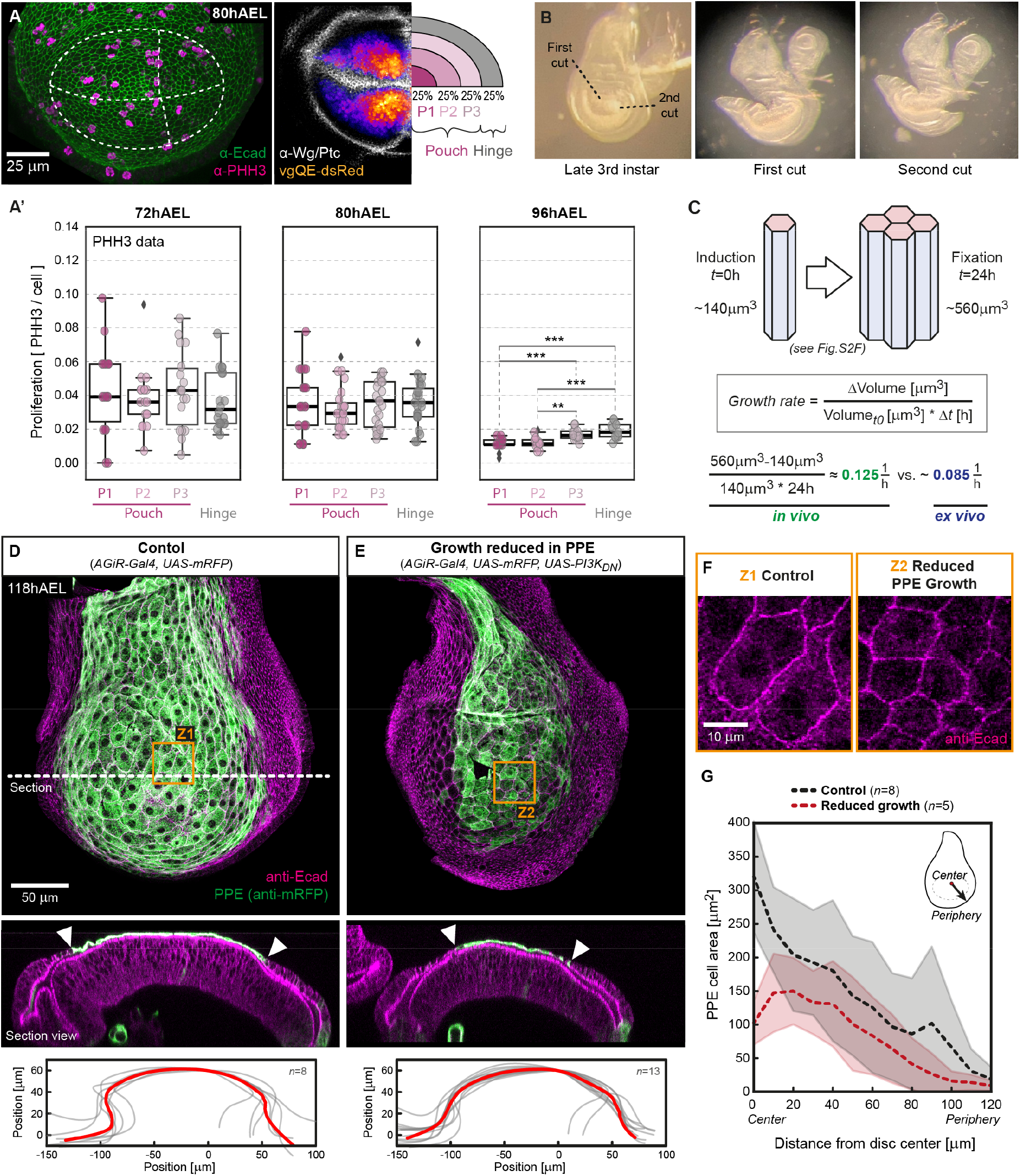
Proliferation pattern and growth modifications in the wing disc. (**A**) *left*: 80hAEL old wing disc stained for E-Cadherin (green) and Phospho-Histone H3 (magenta) to assess proliferation density relative to cell density. The inner Wg ring and the A/P and D/V boundaries are indicated by dotted lines. *right*: Representative 80hAEL wing disc expressing dsRed under control of the *vestigial Quadrant enhancer (vgQE-dsRed*, a marker for the wing pouch) co-labeled for Wg/Ptc. For quantifications the area of the inner Wg ring was subdivided into 4 regions of which the central 3 cover the wing pouch tissue (P1-P3, shades of purple) and the peripheral one corresponds to hinge tissue (gray). (A’) Proliferation density based on PHH3 staining (see panel A), was assessed in individual regions for indicated developmental time points. (B) Wing disc explant before (*left*) and after 2 sequential cuts with performed with a micro-scissor (*middle, right*). The cuts are indicated by dashed lines (*left*). (**C**) *in vivo* clonal proliferation rates can be related to *ex vivo* measured growth rates. The average volume of a DP cell is ~140μm^3^ (see Fig.S2F) at 72hAEL and we know from the clonal essay that in the period between 72 and 96hAEL on average two divisions take place (see Fig.2C’, *right*). From this we can calculate that the average volume of a clone induced at 72hAEL is increasing from 140μm^3^ to ~560μm^3^ within 24h. Hence, we can estimate an *in vivo* growth rate of ~0.125/h. Therefore, the growth rates observed in our *ex vivo* setup are slightly reduced (~0.085/h) compared to *in vivo* growth. Nevertheless, the established *ex vivo* system allows to obtain dynamic volume growth rates in the wing disc and to evaluate potential spatial variation of growth during wing disc development. (**D**) Peripodial projection (*top*) and section view (*middle*) of a 118hAEL wing disc expressing RFP in peripodial cells (*AGiR-Gal4*). In controls, the RFP-marked PPE covers large parts of the disc. The extent of the PPE is marked by two arrowheads in the section view below. Quantifications show these discs form a dome of ~60μm height (bottom) as observed previously (see Fig.1B’). (**E**) Overexpression of a dominant-negative form of PI3K (PI3K_DN_) in peripodial cells results in reduced cellular growth and reduced PPE area (smaller green domain). Despite reduced PPE growth discs still form a dome of similar extent as control discs (*bottom*). (**F**) Magnification of the central PPE region marked by orange rectangles in (D+E). (**F**) Quantification of peripodial apical cell area in control disc (black) and in disc where PPE growth was reduced by PI3K_DN_ (red). In control discs peripodial cell area is maximal in the center of the disc and decreases towards the periphery. In PI3K_DN_ discs the central PPE cell area is significantly reduced. (Error band indicates standard deviation).

**Figure S4.**
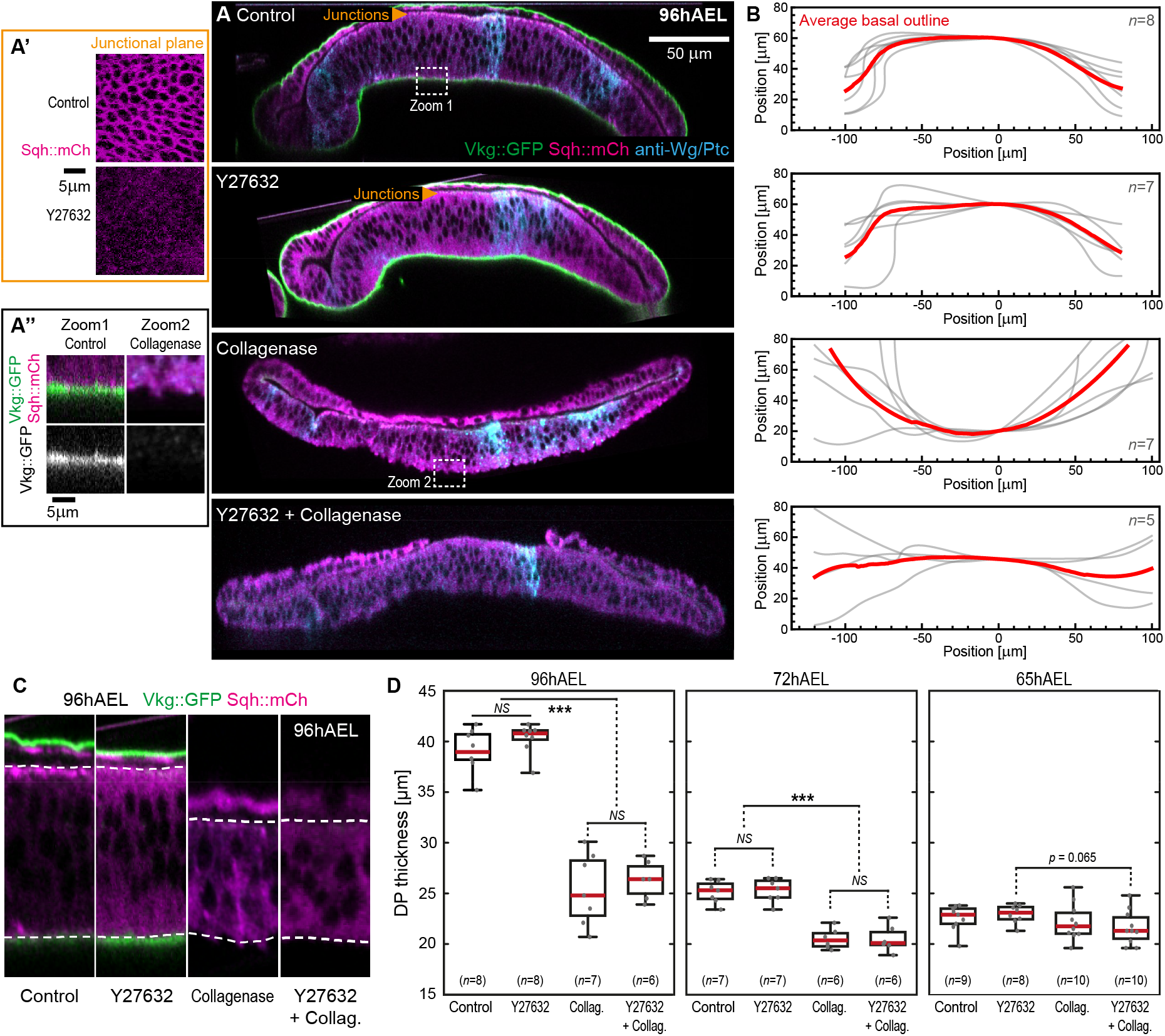
Acute digestion of the ECM concomitant with MyoII inhibition. (**A**) Optical cross-sections parallel to the D/V boundary of representative *Vkg::GFP* (ECM) and *Sqh::mCherry* (MyoII) wing discs at 96hAEL upon the following treatments: *top*: not-treated control, *middle-top*: Rock inhibitor Y27632 treated (MyoII inhibition), *middle-bottom*: Collagenase treated (loss of the ECM) and *bottom*: double treated (Y27632+Collagenase). *Sqh::mCherry* signal in the junctional plane (see orange arrow heads) is observed in-plane-views in control discs (A’ *top*) but lost upon treatment with Y27632 (A’ *bottom*). The ECM labeled by Vkg::GFP (see A’’ zoom1 for control ECM) is completely lost upon treatment with Collagenase (A’’ zoom2). (**B**) Quantifications of the average basal outline of the DP epithelium for the conditions shown in (A). Individual profiles are shown in grey, the average profile in red. While control and Y27632 treated discs maintain domed morphology, the loss of the ECM layer (by Collagenase treatment) results in significant deformation of the epithelial layers: In the presence of MyoII activity (*middle-bottom*) discs tend to inverse their shape and bend upwards. In contrast, a loss of both, MyoII and the ECM results in a nearly flat and relaxed epithelial layer. (**C**) Representative cross sections of the region close to the A/P-D/V intersection of the indicated treatments. The apical and basal surface of the DP epithelium is indicated by dashed lines. (**D**) Quantification of DP epithelial thickness close to the A/P-D/V intersection for different developmental time-points and treatments. While the inhibition of MyoII does not significantly affect DP thickness at any time-point, a loss of the ECM results in a significant reduction in DP thickness at 96 and 72hAEL (in both the Collagenase and the Y27632+Collagenase treated discs).

**Figure S5.**
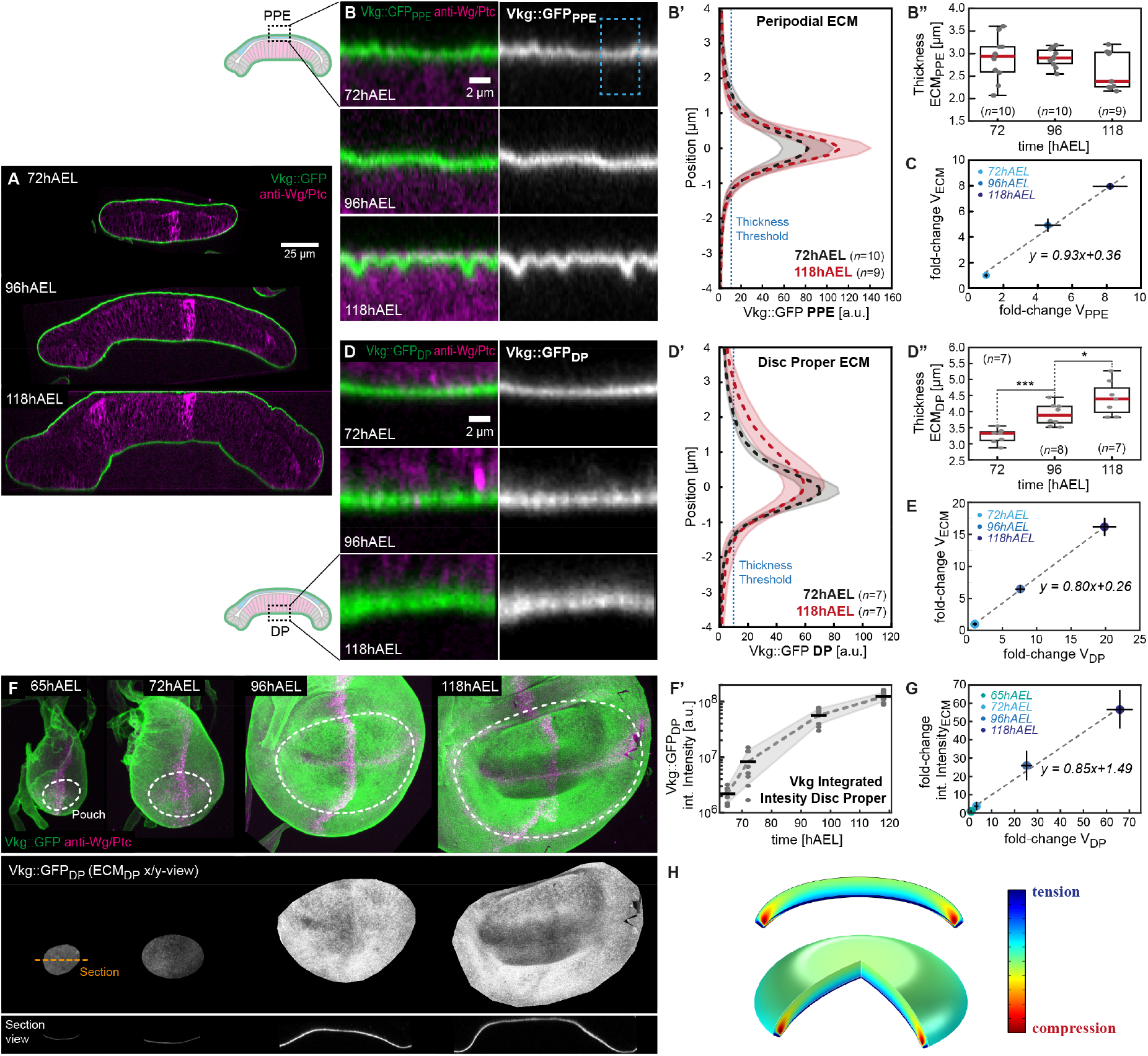
The disc proper epithelium outgrows its ECM layer. (**A**) Cross-sections parallel to the D/V boundary of discs expressing a GFP-tagged version of *Drosophila* Collagen IV (Viking, Vkg::GFP). A shell of ECM (green) can be observed covering the basal side of the epithelial cells at all observed stages. (**B**) Magnifications of representative sections of the peripodial ECM layer (Vkg::GFP_PPE_) at defined stages. Towards the end of development, the ECM_PPE_ tends to be wrinkled. (**B’**) Average plots of Vkg::GFP intensity along the apical-basal axis (as indicated in dashed box in (B)). An increase in Vkg::GFP_PPE_ peak density is observed from 72 to 118hAEL. (**B’’**) However, when ECM_PPE_ thickness is quantified based on an intensity threshold (see dashed blue line in (B’), no change in thickness is observed. See methods for details on intensity profiles and thickness quantifications. (**C**) Plot of relative peripodial epithelial volume increase versus relative top ECM_PPE_ volume increase. Peripodial volume was segmented in 3D (see Fig.S1 H-J and methods) using the inner Wg ring as a landmark. Notably, the top ECM_PPE_ and the peripodial layer show very similar volume growth (slope of linear correlation 0.93). (**D**) Representative section of the bottom ECM_DP_ underlining the basal surface of the DP epithelium. A clear increase in thickness is visible from 72 to 118hAEL. This tendency to increase in thickness can be quantified in Vkg::GFP_DP_ apical-basal intensity profiles (**D’**). A significant increase in thickness is quantified in (**D’’**). (**E**) Plot of relative DP volume changes versus relative estimated ECM volume (obtained by multiplying the area of the inner Wg-ring with the ECM thickness obtained in (D’’)). We observe a linear correlation with a slope of ~0.8. (**F**) *top*: Maximum projections of wing discs expressing Vkg::GFP (green), stained for Wg/Ptc (magenta) at indicated stages. The inner Wg ring, indicating the pouch, is marked by a dashed white line. *middle/bottom*: Using the inner Wg ring we have masked and extracted the Vkg::GFP_DP_ signal within the inner Wg ring underlining the DP. Projections in-plane (*middle*) and cross-section view (*bottom*). (**F’**) The integrated fluorescent intensity of the Vkg::GFP_DP_ underlining the DP (as shown in (F)) should be proportional to Vkg::GFP levels and hence approximates ECM deposition and growth. (**G**) Plot of the relative volume increase of the disc proper epithelium versus the relative increase in integrated Vkg::GFP intensity. DP growth and Vkg::GFP integrated intensity show a linear correlation with a slope of ~0.85. These two observations (E+G) imply a volumetric growth mismatch between the DP epithelium and the underlying ECM_DP_, where the DP outgrows the ECM_DP_ layer. (**H**) Assuming that the ratio of pouch volume increase divided by the ECM_DP_ volume increase between 65h and 118h is 0.79, and that the growth of both layers is planar, we find that this growth mismatch is not sufficient to predict the correct tissue geometry.

**Figure S6.**
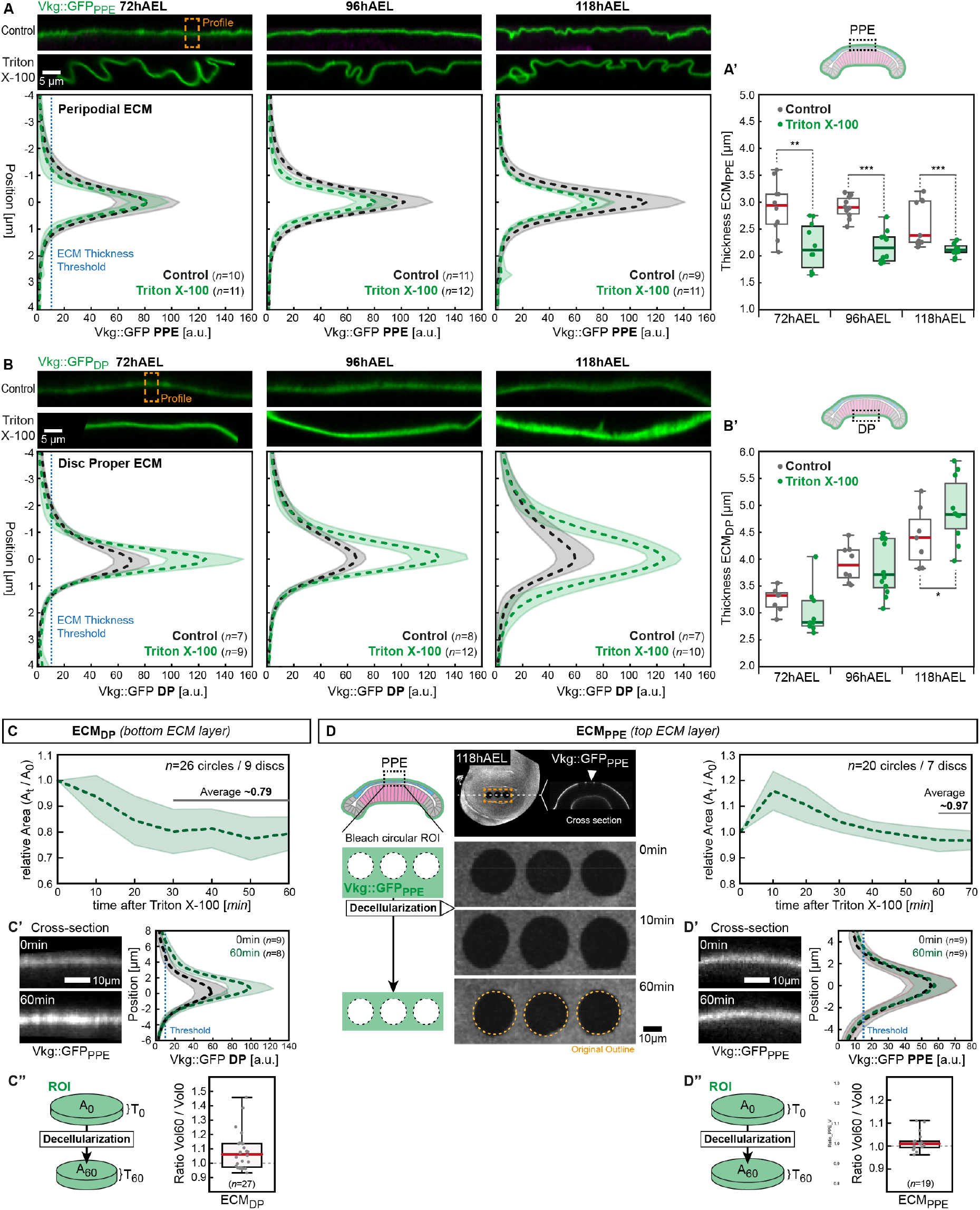
Changes in ECM morphology during larval growth (decellularization) (**A**+**B**) Summary of the changes observed upon decellularization in fixed samples at different developmental stages. The shown data originates from two data-sets acquired under identical imaging conditions (one for the PPE (A) and one for the DP (B)). Hence, fluorescent intensities and profiles are comparable between different timepoints and conditions. (**A**) *top-middle*: Cross-sections of the peripodial ECM labeled by Vkg::GFP in control (*top*) and decellularized (*middle*, Triton X-100 treated) wing discs at indicated age classes. *Bottom*: Profiles of peripodial Vkg::GFP levels (ECM_PPE_) in control (black) and decellularized conditions (green) Error bands indicate standard deviation. (**A’**) Quantification of peripodial ECM thickness at indicated time-points in control (black) and decellularized discs (green). During development we do not observe a significant change in ECM thickness, however, upon decellularization ECM thickness decreases. (**B**) Same as in (A) but for the bottom ECM_DP_. (**B’**) In contrast to the peripodial ECM, the thickness for the bottom ECM_DP_ increases significantly during development. (**C**) Bottom ECM_DP_ relative circular area plotted over time after the addition of Triton X-100. The relative area decreases for 30min before reaching a plateau value at ~79% of the original area. (**C’**) *left*: Cross-section of representative sections of the bottom ECM_DP_ before and 60min after Triton X-100 addition. *right*: Quantification of Vkg::GFP_DP_ intensity before and after decellularization. The profile of the relaxed bottom ECM_DP_ shows increased peak intensity and increased width at the threshold value (dotted blue line, 10a.u.) chosen to quantify ECM thickness in Fig.5D’’. (**C”**) Quantification of changes in estimated circular ECM volume (area *A* * thickness *T*) upon decellularization. (**D**) Results for *ex vivo* decellularization in the ECM_PPE_ layer. *left*: Three circular regions of interest (ROIs) were marked by photobleaching onto the ECM_PPE_ of 118hAEL Vkg::GFP wing discs. *right*: Relative area changes after addition of Triton X-100. In contrast to the bottom ECM_DP_, the relative area in the top ECM_PPE_ transiently increases before reaching a plateau at ~97% of original area after 60-70min. (**D’**) ECM_PPE_ thickness and Vkg::GFP density does not significantly change upon decellularization. (D”) Estimated ECM_PPE_ volume marked by Vkg::GFP remains constant upon decellularization. Error bands show standard deviation.

**Figure S7.**
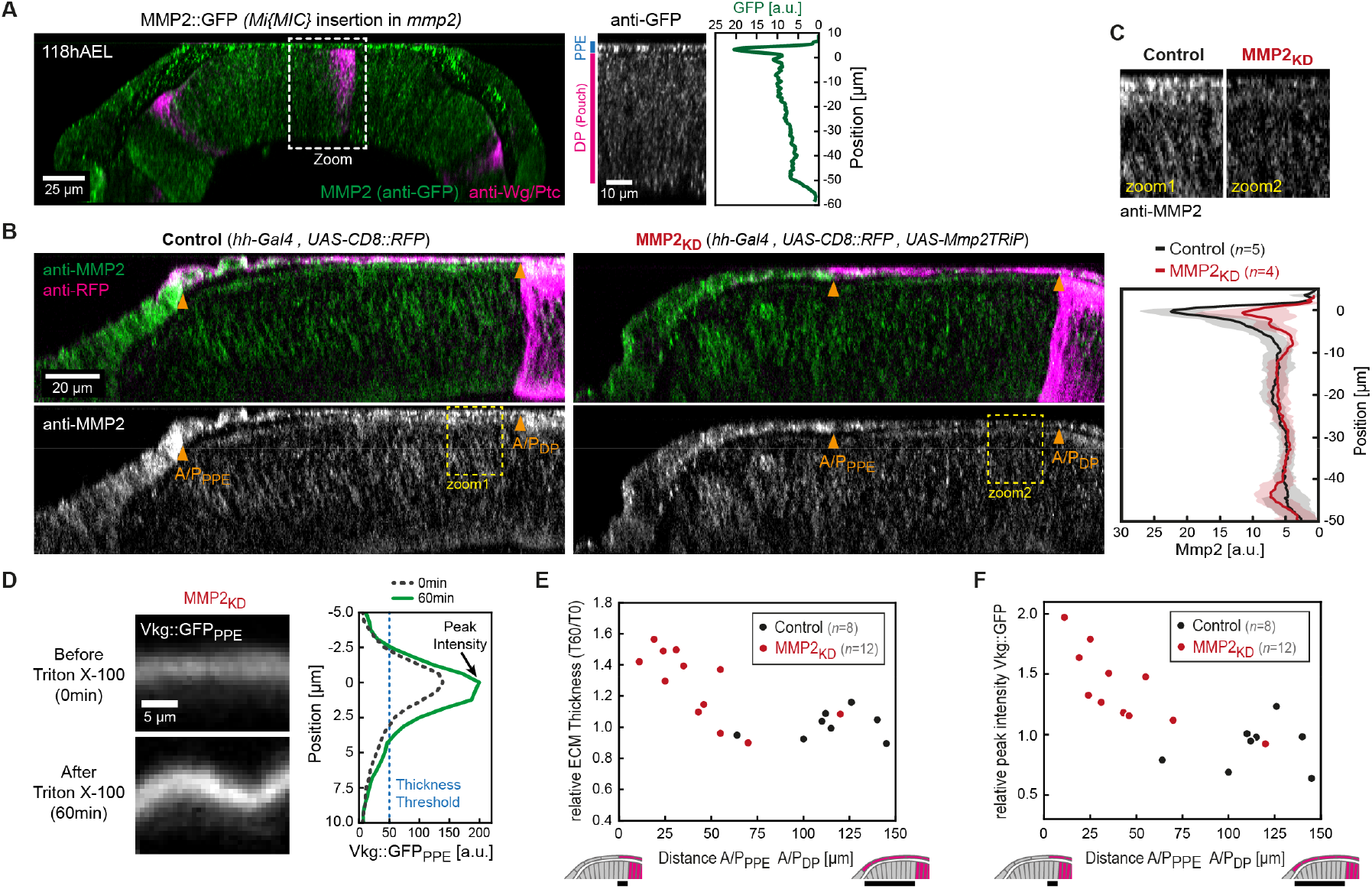
Mmp2 localization and specificity of TRiP-mediated knock-down. (**A**) Section of a wing disc expressing a GFP-tagged form of MMP2 (via Mi{MIC} insertion) at 118hAEL. Consistent with staining for endogenous MMP2, MMP2::GFP is predominantly observed in the PPE. *right*: Magnification of the region indicated by the rectangular box (zoom) and intensity profile of the zoomed region corresponding to MMP2 levels. (**B**) *left*: Control wing disc stained for MMP2 (green) and expressing RFP in the posterior compartment (hh-Gal4). Arrowheads indicate the anterior-posterior (A-P) compartment boundary in the PPE and DP layer. *right*: Knock-down of MMP2 in the posterior compartment results in reduced MMP2 levels in the posterior PPE cells. (**C**) top: Magnifications of the regions marked by yellow rectangles in (B) *Bottom*: Average MMP2 profiles of control (black) and MMP2_KD_ discs (red). The domain between *x*=0 to *x*=5 corresponds to the PPE. In MMP2_KD_ discs MMP2 levels are reduced in the peripodial layer (error bands indicate standard deviation). (**D**) Magnification of peripodial ECM posterior of the A/P boundary before (*top*) and after decellularization (*bottom*). Vkg::GFP intensity profiles were extracted and ECM thickness and peak intensity quantified. Thickness was assessed at a threshold value (50 a.u.). Peak intensity was defined as the maximum intensity of the Vkg::GFP profile. (**E**) Relative change of ECM_PPE_ thickness (Thickness_60min_ / Thickness_0min_) upon decellularization plotted against the distance between the peripodial and disc proper A/P boundary. While in control wing discs this distance is typically ~120μm at the end of 3rd instar development, in MMP2_KD_ this distance decreases with increasing knock-down efficiency. Consistently, the discs showing the strongest ECM thickness changes also show strong reduction in A/P compartment boundary distance. (**F**) Relative Vkg::GFP peak intensity changes (Intensity_60min_ / Intensity_0min_) upon decellularization plotted against A/P boundary distance as in (E).

## Supplementary information

### 1 Description of the multi-layer wing disc model

#### 1.1 Theoretical framework

To gain insight into the shape of the growing wing disc and estimate the growth-induced stress, we constructed a 3D finite element model of an entire wing disc in COMSOL and implemented an existing theory of tissue growth that has been applied successfully to arteries and brain tissue [9, 13, 12].

The tissue grows according to a specified growth deformation tensor, 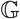, and therefore, the complete deformation gradient 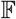 consists of two components: The growth component 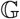 and the elastic component 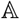, i.e. 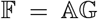. This model is known as morphoelasticity and the multiplicative decomposition is explained in more detail in the caption of Fig. 1. We also made the assumption that the tissue hyperelastic nearly-incompressible neo-Hookean material [12, 7, 8]with a strain-energy density (free energy written in the reference configuration) *W*, given by [1, 9, 6]

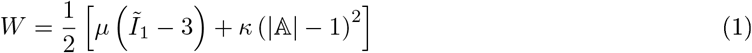

where *μ* and *κ* are the shear and bulk modulus of the material, respectively, and 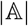 is the determinant of the elastic deformation gradient 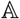. Further, 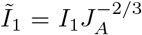, where *I*_1_ is the first invariant of the right Cauchy-Green deformation tensor. Finally, the Cauchy stress tensor is given by

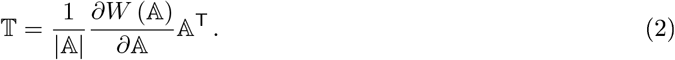

To obtain the deformation of the body with a prescribed growth tensor 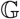, we solve the balance of linear momentum div 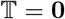. We assume that the external boundary of the wing disk is traction-free, 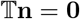, where **n** is the outward facing surface normal. hese are coupled with and the morphoelastic decomposition 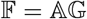, in conjunction with the constitutive law (2).

The full set of boundary conditions is shown in Fig. 2.

In a polar cylindrical basis {**E**_*R*_, **E**_*θ*_, **E**_*Z*_}, we assume that the growth tensor takes the form

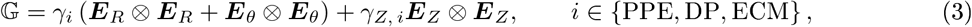

where in the cylindrical plane growth is isotropic if *γ_i_* = 1, and anisotropic of the axial direction is different, *γ_i_* ≠ *γ_Z,i_*. We denote 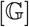 the components of 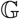 in the polar cylindrical basis:

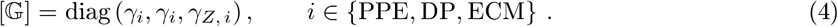

The volume of the wing disc can be computed from the post-grown stress-free configuration 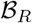, see Fig. 1. This is done by adding the volume of the three discs to which the growth tensor (3) has been applied:

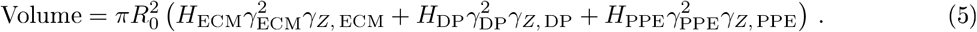

The volume of the deformed configuration will be the same in the limit *κ* → ∞, which corresponds to an incompressible material.

**Figure 1:**
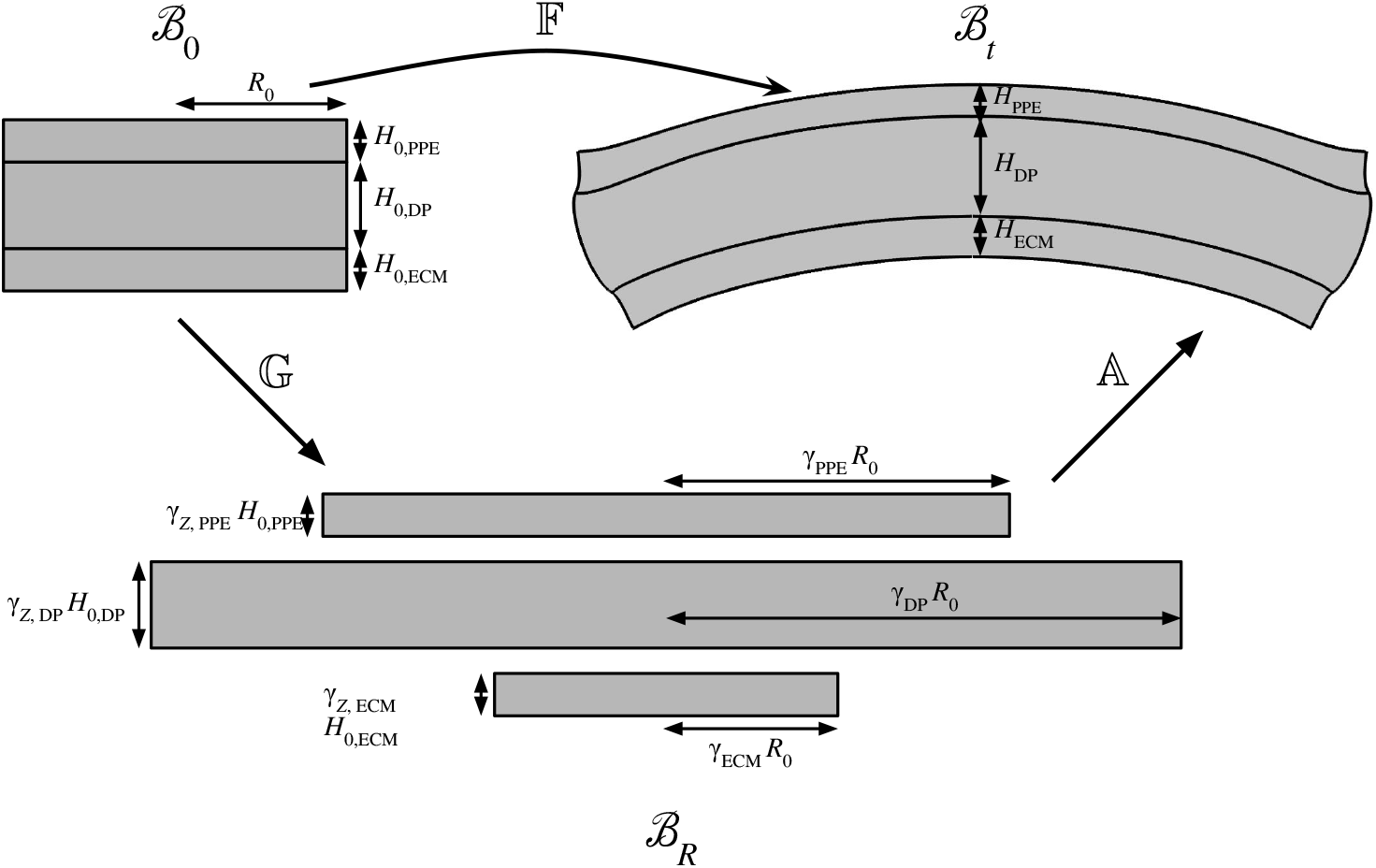
The wing disc is modeled as a multi-layer structure comprised of three layers or fewer. In the initial configuration 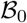, the tissue is ungrown and unstressed. The growth tensor 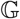 describes growth with out stress, leading to an unstressed incompatible post-grown configuration 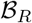, in which the individual layers have growth but do not fit into Euclidean space without breaking the connection between layers. Finally, the elastic deformation gradient 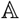 restores compatibility by introducing residual (internal) stress, bringing the body into the current (observed) configuration 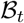. The initial geometry in configuration 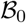 are two glued-together discs of radius *R*_0_. The peripdoial epithelium (PPE) is modeled as a disc of heigth *H*_PPE_, the disc pouch (DP) as a disc of heigth *H*_DP_, and the extracellular matrix (ECM) as a disc with heigth *H*_ECM_. In the post-grown configuration 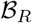, the three discs have radius *γ_Z,i_ R*_0_ and heigth *γ_Z,i_ H_i_* with *i* ∈ {PPE, DP, ECM}, where the growth tensor in the polar cylindrical basis is 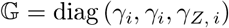.

#### 1.2 Four different scenarios

##### 1.2.1 Scenario: Non-uniform growth of one single layer

In this scenario, we assume that the wing disc is made up of only one layer, the disc pouch (DP). We explore the possibility of inducing a deformation of the disc through non-uniform growth pattern in the disc plane. The peripodial epithelium (PPE) as well as the extracellular matrix (ECM) are not modelled, as reflected in the parameter overview, see Table 1 column “inner vs outer”. We model the uniformity by breaking the domain into an inner ring and an outer ring. We denote the radial variable and the disc radius in the initial configuration 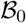 by *R* and *R*_0_, respectively. Then the inner region shall be defined by *R* ≤ *R*_0_/2 and the outer region by *R*_0_/2 ≤ *R* < *R*_0_. In the inner and outer regions, we prescribe growth tensors 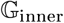 and 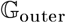, respectively:

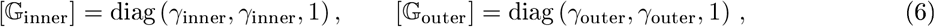

where for the sake of simplicity we assumed no growth in axial direction (*γ*_*Z*,inner_ = *γ*_*Z*, outer_ = 1)· The results of simulations with increasing ratio of *γ*_inner_/*γ*_outer_ are shown in Fig. 2B. An increasing non-uniformity *γ*_inner_/*γ*_outer_ creates a swollen inner region, but the lack of boundary constraints and the softness of the material given realistic geometric and stiffness parameters (see Table 1) excludes a doming of the disc as observed in the wild type.

##### 1.2.2 Scenario: Bilayer PPE-DP

In this scenario, we test the hypothesis that the wing disc consists of two layers, a fast growing peripodial epithelium and a slower growing wing pouch. The ECM is not modeled in this scenario, see Table 1 column “PPE vs DP”. For the sake of simplicity, we consider planar growth in both PPE and DP, that is

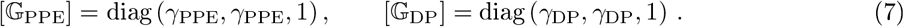

The results of simulations with *γ*_PPE_ = 4.3 and *γ*_DP_ = 3.31, which corresponds to a volume ratio between PPE and DP layers of 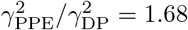, are shown in Fig. 3B. However, as discussed in the main text, experiments show that growth between PPE and DP is nearly compatible, demonstrate that epithelial doming and thickening are not due to a non-uniformity of growth within or between epithelial layers.

**Figure 2:**
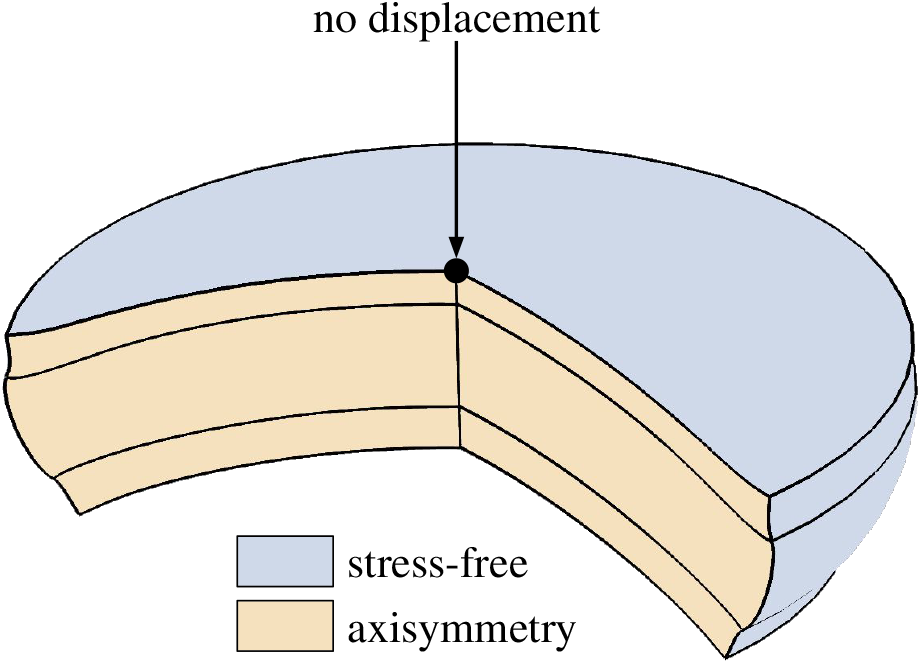
Boundary conditions for *Drosophila*, wing disc sandwich model. The disc is assumed axisymmetric, with a no-stress boundary condition ***Tn*** = 0 imposed at the surface. The no-displacement boundary condition on the symmetry axis (black dot) serves to eliminate translational degrees of freedom for the `Comsol` solver.

##### 1.2.3 Scenario: Bilayer DP-ECM, with differential growth anisotropy

In the main text, we describe how a two layer system made up of a DP layer and ECM layer capures the bending of the wing disc, as well as a number other of experimentally measured geometric quantities, see Fig. 6. In particular, the DP layer grows in plane whereas the ECM layer deviates from planar growth.

The growth tensors describing the two layers are

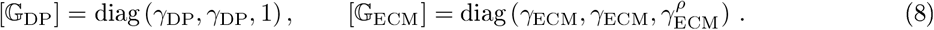

The components of the growth tensor for the DP satisfy *γ*_DP_ = 1 at *t* = 65h and 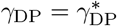 at *t* = 118h and for the ECM they satisfy *γ*_ECM_ = 1 at *t* = 65h and 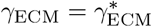 at *t* = 118h.

For the disc pouch, the parameter 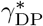 can be determined by fitting the linear function

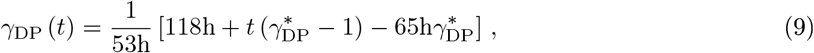

which satisfies the above constraints. Since the volume over time in the DP, *V*_DP_ (*t*), is available from experiments, we now show how *γ*_DP_ (*t*) can be related to experimental volume measurements. The volume of the DP is given by 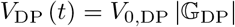, where |·| = det (·) denotes the determinant of a tensor. Here, 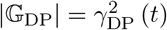. The last two equations can be combined to

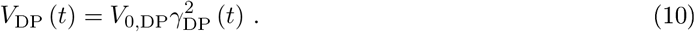

The initial volume *V*_0,DP_ at *t* = 65h can be computed as the volume of a cylinder, 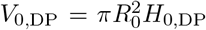 where from experimental measurements we used values for the initial radius, *R*_0_, and initial heigth of the DP, *H*_0,DP_ as stated in Table 1 column “best fit DP vs ECM” (see the initial state, 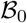, in Fig. 1). With this information, we can solve (10) for 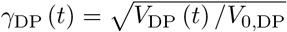. We make a least squares fit, using the experimentally measured values *V*_DP_ (*t*) and *V*_0,DP_ and using the form of (9) for *γ*_DP_ (*t*). As a result of the least squares fit, we obtain 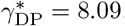.

To obtain 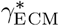, we proceed similarly. We use a linear functional form for *γ*_ECM_(*t*), which is identical to (9) (with the subscript DP replaced by ECM). The ECM volume, on the other hand, depends on *ρ*, since 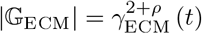. Thus, the ECM volume is given by

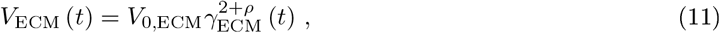

where 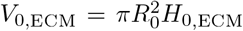 and values for *R*_0_, *H*_0,ECM_ are stated in Table 1 column “best fit DP vs ECM”. With this information, we can solve (11) for 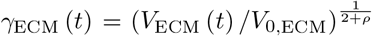. So for a given value of *ρ*, we make a least squares fit from experimental values *V*_ECM_ (*t*) and *V*_0,ECM_ and using the linear form for *γ*_ECM_ (*t*) given above. We repeat the least squares fitting for a series of values of *ρ*, obtaining data points 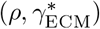 from the best fits, see Fig. 3. This data is well described by the smooth function 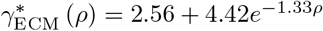.

**Figure 3:**
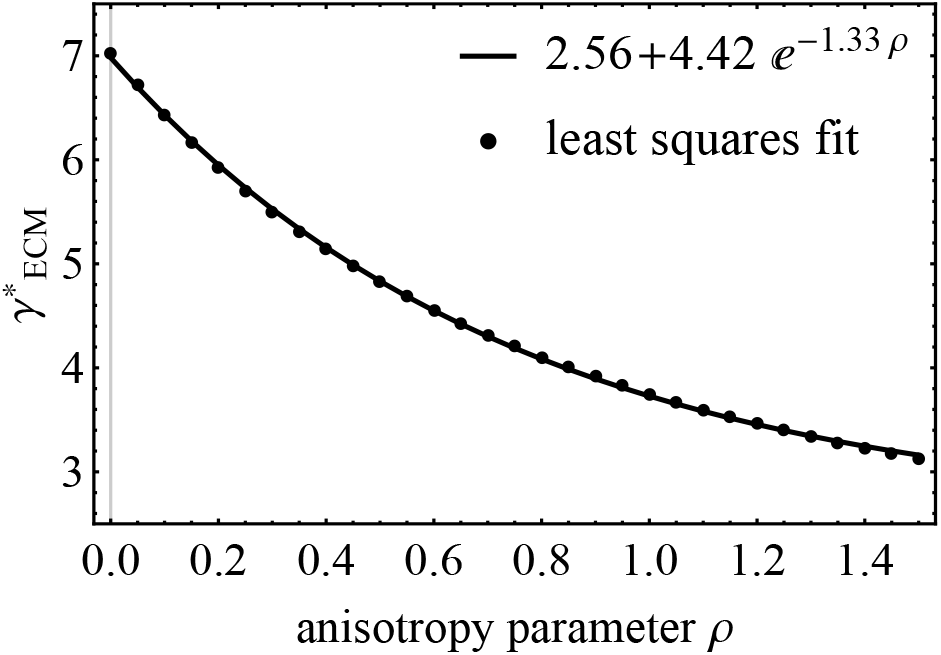
Obtaining a relationship between 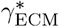 and *ρ*, using volumetric data of the ECM. In the inplane scenario *ρ* = 0, all of the volume increase in the ECM is assumed to be distributed in the plane, and the in-plane growth component 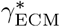 is highest in this case. As *ρ* increases, an increased amount of the measured volume is assumed to be distributed in *Z*-direction, so that the in-plane growth component 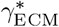 decreases. The special case *ρ* =1 represents isotropic growth, in which case the ECM growth tensor takes the form 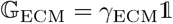 where 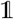 is the 3-dimensional identity.

With the parameters 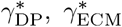 determined from volume data, the remaining parameters *ρ* and *μ* = *μ*_DP_/*μ*_ECM_ are determined in Fig. 6B. There, we consider three regions which show where the relative error between the simulated values is within a certain tolerance of the the mean of experimental values, measured at the end of the observed window at *t* = 118h. The rose region compares the measured and simulated disc pouch thickness *H*_DP_, the blue region the measured and simulated ECM thickness *H*_ECM_. Finally, the orange region compares the ratio of reference thickness to observed thickness, *H*_*r*,ECM_/*H*_ECM_, where we denoted the reference configuration with the subscript *r*. Since 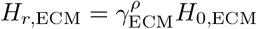. Thus, the orange region compares the measured and simulated quantity 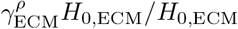.

When all three quantities, that is pouch thickness *H*_DP_, ECM thickness *H*_ECM_ and relative thickness increase upon decellularization 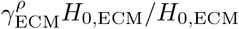 are taken together, we obtain the region diagram in Fig. 6B. When all three quantities are simultaneously below the relative error tolerances specified inside the figure, we obtain the dark region. This region allows us to determine the best fit *ρ* = 0.45 and *μ* = 25.

##### 1.2.4 Scenario: Bilayer DP-ECM, both layers growing in-plane

This scenario serves as a demonstration that differential growth anisotropy is indeed essential to capture the wing disc morphology. In Fig. S5H, using the parameter values given in Table 1 column “in-plane DP vs ECM”, we explore the scenario from Section 1.2.3 but with a crucial difference: There is no differential growth anisotropy (*ρ* = 0), meaning that both the DP and ECM grow in-plane. The result is shown in Fig. S5H, demonstrating clearly that without growth anisotropy, the correct wing disc morphology can not be achieved, we end up with a structure that is far too flat.

### 2 Numerical implementation

Numerical simulations are obtained using a finite clement code that solves the equation of finite elasticity on a multi-layer structure of glucd-togcthcr axisymmctric cylinders. The 2D axisymmetric numerical problem is solved in Comsol Multiphysics ^®^ [3]. In Comsol, the morphoelatic decomposition 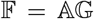 cannot be directly entered into the software. We give here a succinct summary of the implementation procedure stated in works of Larry Taber [7, 12], showing how growth problems can be studied in Comsol.

**Table 1:** Simulation parameter values for the different scenarios described in Sec. 1.2.

|  |  | inner vs outer<br>(Fig. 2B) | PPE vs DP<br>(Fig. 3) | best fit DP vs<br>ECM (Fig. 6) | in-plane DP vs<br>ECM (Fig. S5H) | reference |
| --- | --- | --- | --- | --- | --- | --- |
| geometric | $R_0$ [$\mu\text{m}$] | 20.05 | 20.05 | 20.05 | 20.05 | measured |
| | $H_{0,\text{PPE}}$ [$\mu\text{m}$] | - | 5.25 | - | - | measured |
| | $H_{0,\text{DP}}$ [$\mu\text{m}$] | 22.36 | 22.36 | 22.36 | 22.36 | measured |
| | $H_{0,\text{ECM}}$ [$\mu\text{m}$] | - | - | 3.09 | 3.09 | measured |
| stiffness | $\mu_{\text{PPE}}$ [kPa] | - | 1 | - | - | refs [11, 4, 10] |
| | $\mu_{\text{DP}}$ [kPa] | 1 | 1 | 1 | 1 | |
| | $\mu_{\text{ECM}}$ [kPa] | - | - | 25 | 25 | Fig. 6B |
| growth at 118h | $\gamma_{\text{PPE}}$ [1] | - | 3 | - | - | - |
| | $\gamma_{\text{DP}}$ [1] | 1-2 | 2.5 | 8.09 | 8.09 | volume data |
| | $\gamma_{\text{ECM}}$ [1] | - | - | 4.99 | 6.99 | volume data |
| | $\rho$ | - | - | 0.45 | 0 | Fig. 6B |

#### 2.1 General case

Comsol computes derivatives with respect to displacement gradients to obtain a second Piola-Kirchhoff stress tensor

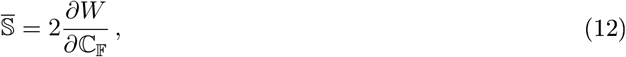

where we denote 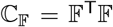 the right Cauchy-Green strain tensor of the total deformation gradient 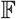. In particular, the finite element formulation requires the stress 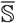 per unit initial area 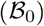. The second Piola-Kirchhoff stress in terms of Cauchy stress is given by

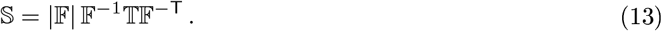

For an incompressible or nearly incompressible material, the Cauchy stress is given by (2), which can be rewritten as

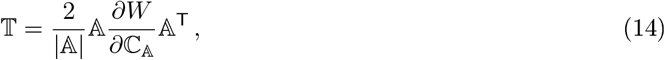

where we denote 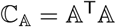 the right Cauchy-Green strain tensor of the elastic deformation gradient 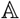. Inserting (14) into (13), we get

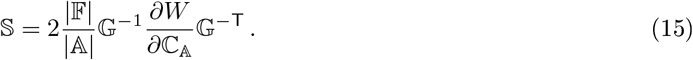

As shown in [7], the gradient of *W* satisfies the following transformation relationship:

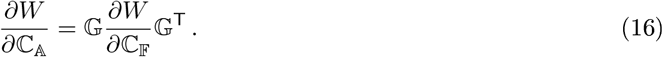

Inserting this into (15), we find

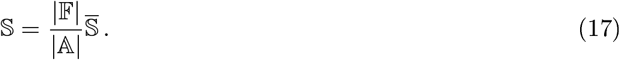

So the appropriate expression of the second Piola-Kirchhoff stress can be obtained by multiplying the equation for 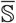, which Comsol uses by default, by 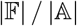. In addition, *W* must be defined in Comsol in terms of the components of 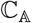, that is

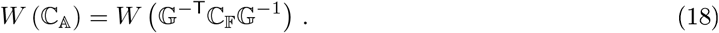

#### 2.2 Specific case

Note that Comsol will solve for 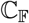. Therefore, it is necessary to provide Comsol with the components of 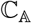 in terms of the components of 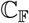, in order for a successful implementation of the modified strain energy *W* and and the modification (17). In our problem, we assume that the growth tensor is of the structure olar cylindrical basis {**E**_*R*_, **E**_*θ*_, **E**_*Z*_} and has diagonal form

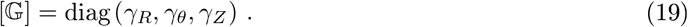

Furthermore, the components 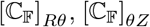 are fixed by the constraint of axisymmetry. Further recalling that 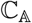 is symmetric, the four unique components 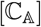 are for this particular geometry:

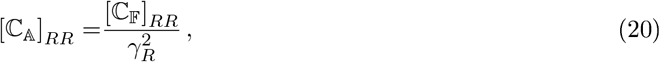

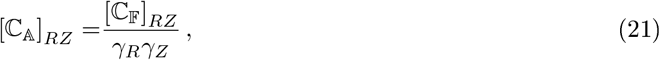

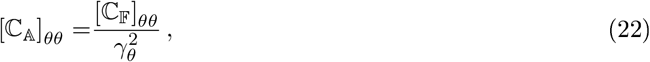

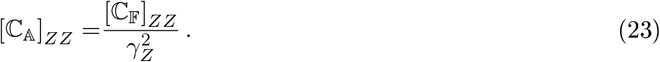

These can be inserted into the re-defined strain energy density for Comsol, (18), and used to compute the determinant 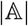 for the re-defined second Piola Kirchhoff stress (17).

### 3 Validation of the numerical implementation against exact solution

In this section, we test the computational framework presented in Section 2 by comparing it with an analytically known solution for an incompressible neo-Hookean material with growth. The setup we are considering is shown in Fig. 4A. We consider a cylinder with an initial radius *R*_0_, with no displacement allowed in *Z*-direction, that is 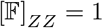. In the analytical axisymmstric calculation, we impose 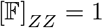 everywhere in the bulk, whereas in the numerical implementation, as shown in Fig. 4A, we impose this constraint on the top and bottom highlighted planes. Further, we impose a growth anisotropy. Our goal is to compare the numerical and analytical expressions for the Cauchy stress tensor and to determine the relative error between them, so that we establish a benchmark for what kind of error to expect in the simulations presented in discussed in Section 1.2 where no analytical solution exists.

#### 3.1 Derivation of exact solution

We consider the case of a single incompressible growing neo-Hookean disk. We assume that there is no deformation at the point of symmetry, and that there are no external forces, so that any deformation is caused purely by growth and the elastic response. This calculation follows a similar path to [5, 2, 6]. For an incompressible disk in the polar cylindrical basis {**E**_*R*_, **E**_*θ*_, **E**_*Z*_}, the morphoelastic decomposition 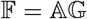 reads

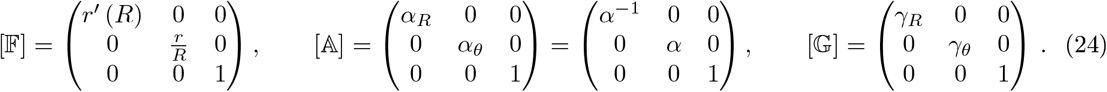

Eliminating *α*, we obtain the kinematic relationship

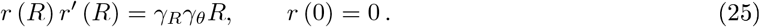

Let *W*(*α_R_*, *α_θ_*) be the strain-energy density, which relates to the Cauchy stress tensor by

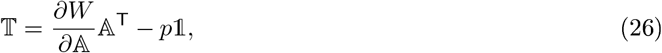

where *p* is the Lagrange multiplier enforcing incompressibility. In components this reads

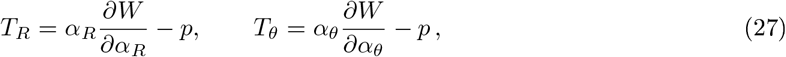

where we used the shorthand notation 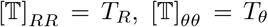. With no external loads, mechanical equilibrium requires div 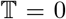, which takes the form 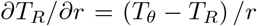. Expressing with respect to the reference radius *R*, this becomes

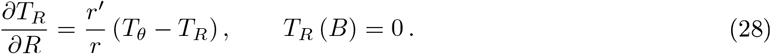

**Figure 4:**
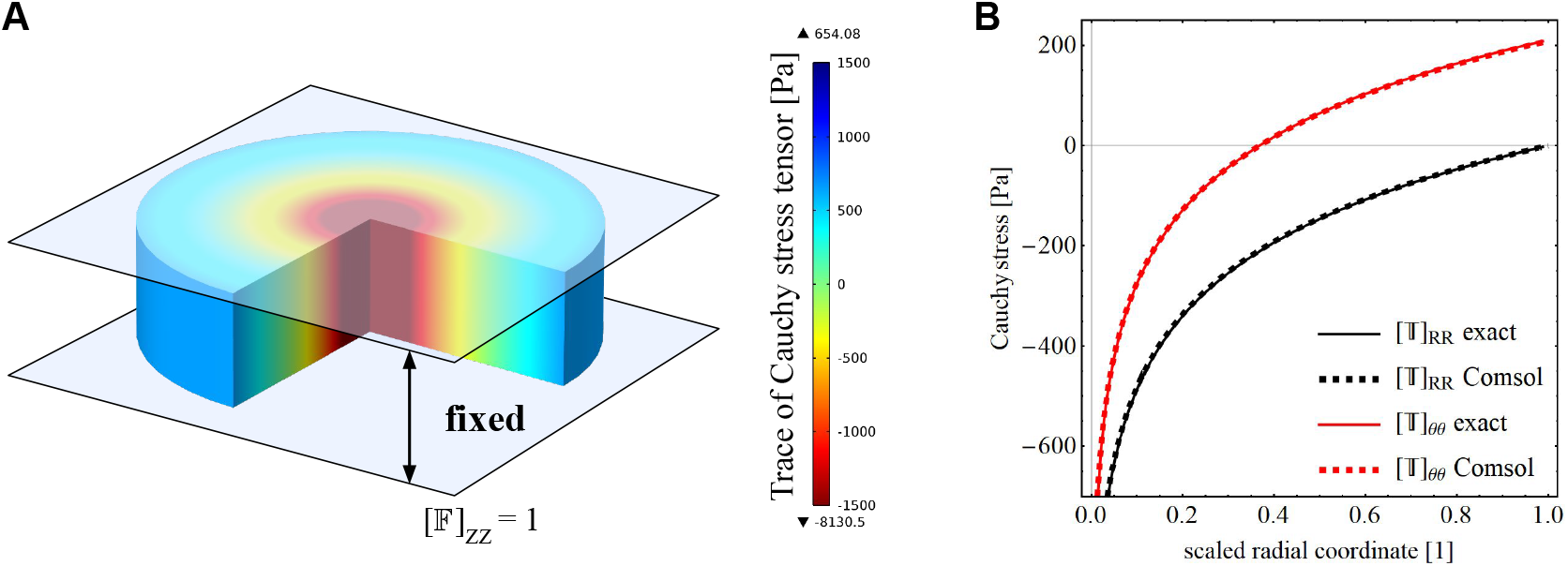
Comparison between analytical solution and numerical implementation in Comsol.

Defining 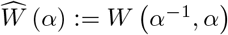, we have

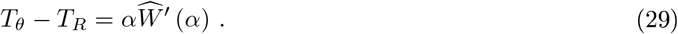

Now taking into account (29), for the radial stress we must solve

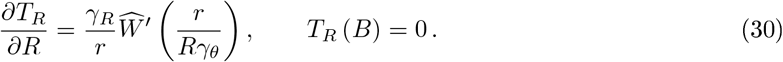

For a neo-Hookean strain-energy density

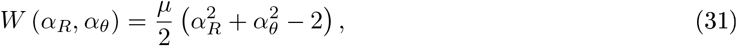

and taking into account the relationships from the morphoelastic decomposition (1) and (25), we can express (30) as

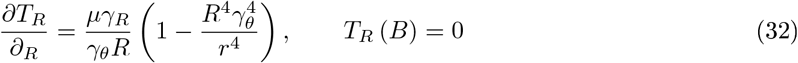

Notice the incorrect factor 2 in Eq. (15) of the refrence [5]. Once the radial stress is known, the circumferential stress can be obtained from (29)

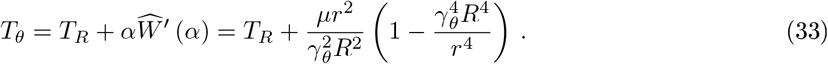

Once again, notice the incorrect factor 2 in Eq. (16) of the refrence [5]. To obtain the full solution, one must provide *γ_R_* (*R*) and *γ_θ_* (*R*) and can then solve the two non-linear coupled ODEs (29) and (32). For *γ_R_*, *γ_θ_* spatially constant, we obtain the exact analytical solution

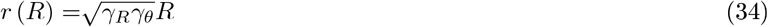

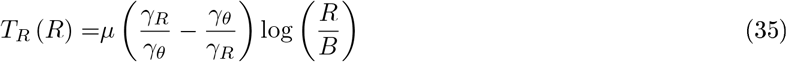

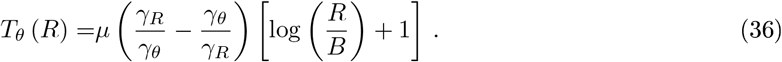

#### 3.2 Comparison of exact and analytical solutions

In Fig. 4, we compare the exact analytical solution (35), (36) with the numerical implementation in Comsol Multiphysics ^®^ as described in Section 2. We obtain excellent agreement, with the largest relative error between numerics and analytics at the disk center being 3.36% and the larges error at the disk edge being 1.38% Given that at the’ disk center, the stress tensor has a singularity (*T* ~ log *R*) as see in (35) and (36), expect the lower error at the edge to be more representative. The FEM mesh in this case was composed of 26006 triangular elements (all simulations presented in Section 1.2 had around 25k triangular elements, always using the Comsol predefined setting “Extremely fine” for the mesh, without any manually defined mesh refinement). Overall, the comparison between exact and analytical solutions shows that the numerical implementation is reliable, and justifying its use for more complex geometries like the sandwich structures presented in the paper.

